# Report of Partial findings from the National Toxicology Program Carcinogenesis Studies of Cell Phone Radiofrequency Radiation in Hsd: Sprague Dawley^®^ SD rats (Whole Body Exposures)

**DOI:** 10.1101/055699

**Authors:** Michael Wyde, Mark Cesta, Chad Blystone, Susan Elmore, Paul Foster, Michelle Hooth, Grace Kissling, David Malarkey, Robert Sills, Matthew Stout, Nigel Walker, Kristine Witt, Mary Wolfe, John Bucher

## Abstract

The U.S. National Toxicology Program (NTP) has carried out extensive rodent toxicology and carcinogenesis studies of radiofrequency radiation (RFR) at frequencies and modulations used in the U.S. telecommunications industry. This report presents partial findings from these studies. The occurrences of two tumor types in male Harlan Sprague Dawley rats exposed to RFR, malignant gliomas in the brain and schwannomas of the heart, were considered of particular interest and are the subject of this report. The findings in this report were reviewed by expert peer reviewers selected by the NTP and National Institutes of Health (NIH). These reviews and responses to comments are included as appendices to this report, and revisions to the current document have incorporated and addressed these comments. When the studies are completed, they will undergo additional peer review before publication in full as part of the NTP’s Toxicology and Carcinogenesis Technical Reports Series. No portion of this work has been submitted for publication in a scientific journal. Supplemental information in the form of four additional manuscripts has or will soon be submitted for publication. These manuscripts describe in detail the designs and performance of the RFR exposure system, the dosimetry of RFR exposures in rats and mice, the results to a series of pilot studies establishing the ability of the animals to thermoregulate during RFR exposures, and studies of DNA damage. (1) Capstick M, Kuster N, Kühn S, Berdinas-Torres V, Wilson P, Ladbury J, Koepke G, McCormick D, Gauger J, and Melnick R. A radio frequency radiation reverberation chamber exposure system for rodents; (2) Yijian G, Capstick M, McCormick D, Gauger J, Horn T, Wilson P, Melnick RL, and Kuster N. Life time dosimetric assessment for mice and rats exposed to cell phone radiation; (3) Wyde ME, Horn TL, Capstick M, Ladbury J, Koepke G, Wilson P, Stout MD, Kuster N, Melnick R, Bucher JR, and McCormick D. Pilot studies of the National Toxicology Program’s cell phone radiofrequency radiation reverberation chamber exposure system; (4) Smith-Roe SL, Wyde ME, Stout MD, Winters J, Hobbs CA, Shepard KG, Green A, Kissling GE, Tice RR, Bucher JR, and Witt KL. Evaluation of the genotoxicity of cell phone radiofrequency radiation in male and female rats and mice following subchronic exposure.

## SUMMARY

The purpose of this communication is to report partial findings from a series of radiofrequency radiation (RFR) cancer studies in rats performed under the auspices of the U.S. National Toxicology Program (NTP).^1^ This report contains peer-reviewed, neoplastic and hyperplastic findings only in the brain and heart of Hsd:Sprague Dawley^®^ SD^®^ (HSD) rats exposed to RFR starting *in utero* and continuing throughout their lifetimes. These studies found low incidences of malignant gliomas in the brain and schwannomas in the heart of male rats exposed to RFR of the two types [Code Division Multiple Access (CDMA) and Global System for Mobile Communications (GSM)] currently used in U.S. wireless networks. Potentially preneoplastic lesions were also observed in the brain and heart of male rats exposed to RFR.

The review of partial study data in this report has been prompted by several factors. Given the widespread global usage of mobile communications among users of all ages, even a very small increase in the incidence of disease resulting from exposure to RFR could have broad implications for public health. There is a high level of public and media interest regarding the safety of cell phone RFR and the specific results of these NTP studies.

Lastly, the tumors in the brain and heart observed at low incidence in male rats exposed to GSM-and CDMA-modulated cell phone RFR in this study are of a type similar to tumors observed in some epidemiology studies of cell phone use. These findings appear to support the International Agency for Research on Cancer (IARC) conclusions regarding the possible carcinogenic potential of RFR.^2^

It is important to note that this document reviews only the findings from the brain and heart and is not a complete report of all findings from the NTP’s studies. Additional data from these studies in Hsd:Sprague Dawley^®^ SD^®^ (Harlan) rats and similar studies conducted in B6C3F1/N mice are currently under evaluation and will be reported together with the current findings in two forthcoming NTP Technical Reports.

## STUDY RATIONALE

Cell phones and other commonly used wireless communication devices transmit information via non-ionizing radiofrequency radiation (RFR). In 2013, IARC classified RFR as a *possible human carcinogen* based on “limited evidence” of an association between exposure to RFR from heavy wireless phone use and glioma and acoustic neuroma (vestibular schwannoma) in human epidemiology studies, and “limited evidence” for the carcinogenicity of RFR in experimental animals. While ionizing radiation is a well-accepted human carcinogen, theoretical arguments have been raised against the possibility that non-ionizing radiation could induce tumors (discussed in IARC, 2013). Given the extremely large number of people who use wireless communication devices, even a very small increase in the incidence of disease resulting from exposure to the RFR generated by those devices could have broad implications for public health.

## DESCRIPTION OF THE NTP CELL PHONE RFR PROGRAM

RFR emitted by wireless communication devices, especially cell phones, was nominated to the NTP for toxicology and carcinogenicity testing by the U.S. Food and Drug Administration (FDA). After careful and extensive evaluation of the published literature and experimental efforts already underway at that time, the NTP concluded that additional studies were warranted to more clearly define any potential health hazard to the U.S. population. Due to the technical complexity of such studies, NTP staff worked closely with RFR experts from the National Institute of Standards and Technology (NIST). With support from NTP, engineers at NIST evaluated various types of RFR exposure systems and demonstrated the feasibility of using a specially designed exposure system (reverberation chambers), which resolved the inherent limitations identified in existing systems.

In general, NTP chronic toxicity/carcinogenicity studies expose laboratory rodents to a test article for up to 2 years and are designed to determine the potential for the agent tested to be hazardous and/or carcinogenic to humans.^3^ For cell phone RFR, a program of study was designed to evaluate potential, long-term health effects of whole-body exposures. These studies were conducted in three phases: (1) a series of pilot studies to establish field strengths that do not raise body temperature, (2) 28-day toxicology studies in rodents exposed to various low-level field strengths, and (3) chronic toxicology and carcinogenicity studies. The studies were carried out under contract at IIT Research Institute (IITRI) in Chicago, IL following Good Laboratory Practices (GLP). These studies were conducted in rats and mice using a reverberation chamber exposure system with two signal modulations [Code Division Multiple Access (CDMA) and Global System for Mobile Communications (GSM)] at two frequencies (900 MHz for rats and 1900 MHz for mice), the modulations and frequency bands that are primarily used in the United States.

## STUDY DESIGN

Hsd:Sprague Dawley^®^ SD^®^ (Harlan) rats were housed in custom-designed reverberation chambers and exposed to cell phone RFR. Experimentally generated 900 MHz RF fields with either GSM or CDMA modulation were continuously monitored in real-time during all exposure periods via RF sensors located in each exposure chamber that recorded RF field strength (V/m). Animal exposure levels are reported as whole-body specific absorption rate (SAR), a biological measure of exposure based on the deposition of RF energy into an absorbing organism or tissue. SAR is defined as the energy (watts) absorbed per mass of tissue (kilograms). Rats were exposed to GSM-or CDMA-modulated RFR at 900 MHz with whole-body SAR exposures of 0, 1.5, 3, or 6 W/kg. RFR field strengths were frequently adjusted based on changes in body weight to maintain desired SAR levels.

Exposures to RFR were initiated *in utero* beginning with the exposure of pregnant dams (approximately 11-14 weeks of age) on Gestation Day (GD) 5 and continuing throughout gestation. After birth, dams and pups were exposed in the same cage through weaning on postnatal day (PND) 21, at which point the dams were removed and exposure of 90 pups per sex per group was continued for up to 106 weeks. Pups remained group-housed from PND 21 until they were individually housed on PND 35. Control and treatment groups were populated with no more than 3 pups per sex per litter. All RF exposures were conducted over a period of approximately 18 hours using a continuous cycle of 10 minutes on (exposed) and 10 minutes off (not exposed), for a total daily exposure time of approximately 9 hours a day, 7 days/week. A single, common group of unexposed animals of each sex served as controls for both RFR modulations. These control rats were housed in identical reverberation chambers with no RF signal generation. Each chamber was maintained on a 12-hour light/dark cycle, within a temperature range of 72 ± 3°F, a humidity range of 50 ± 15%, and with at least 10 air changes per hour. Throughout the studies, all animals were provided *ad libitum* access to feed and water.

## RESULTS

In pregnant rats exposed to 900 MHz GSM-or CDMA-modulated RFR, no exposure-related effects were observed on the percent of dams littering, litter size, or sex distribution of pups. Small, exposure-level-dependent reductions (up to 7%) in body weights compared to controls were observed throughout gestation and lactation in dams exposed to GSM-or CDMA-modulated RFR. In the offspring, litter weights tended to be lower (up to 9%) in GSM and CDMA RFR-exposed groups compared to controls. Early in the lactation phase, body weights of male and female pups were lower in the GSM-modulated (8%) and CDMA-modulated (15%) RFR groups at 6 W/kg compared to controls. These weight differences in the offspring for both GSM and CDMA exposures tended to lessen (6% and 10%, respectively) as lactation progressed. Throughout the remainder of the chronic study, no RFR exposure-related effects on body weights were observed in male and female rats exposed to RFR, regardless of modulation (Figures 1 and 2). At the end of the 2-year study, survival was lower in the control group of males than in all groups of male rats exposed to GSM-modulated RFR (Figure 3).

**Figure 1.**
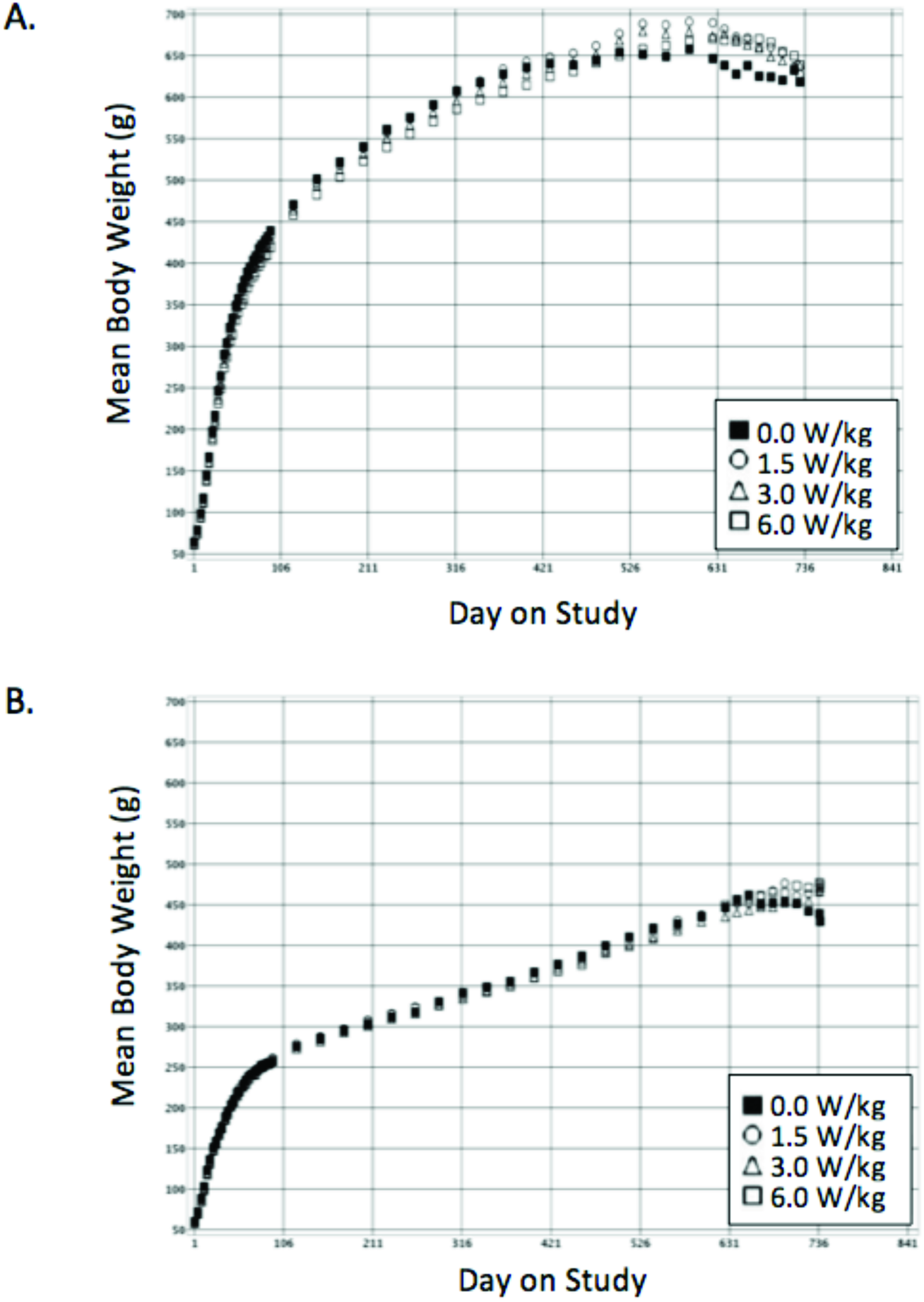
Growth Curves for Male (A) and Female (B) Rats Exposed to Whole Body GSM-Modulated RFR for 2 Years

**Figure 2.**
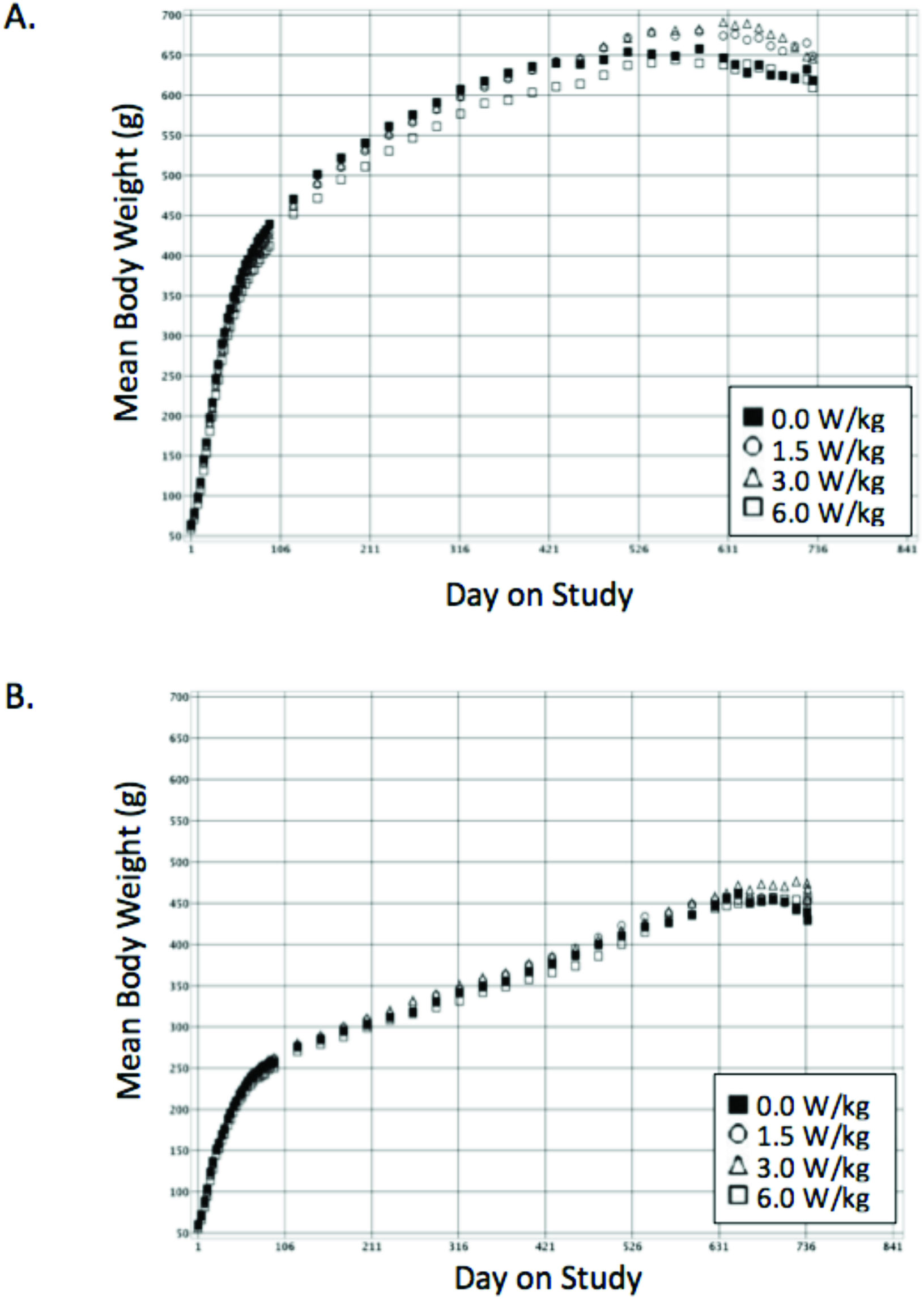
Growth Curves for Male (A) and Female (B) Rats Exposed to Whole Body CDMA-Modulated RFR for 2 Years

**Figure 3.**
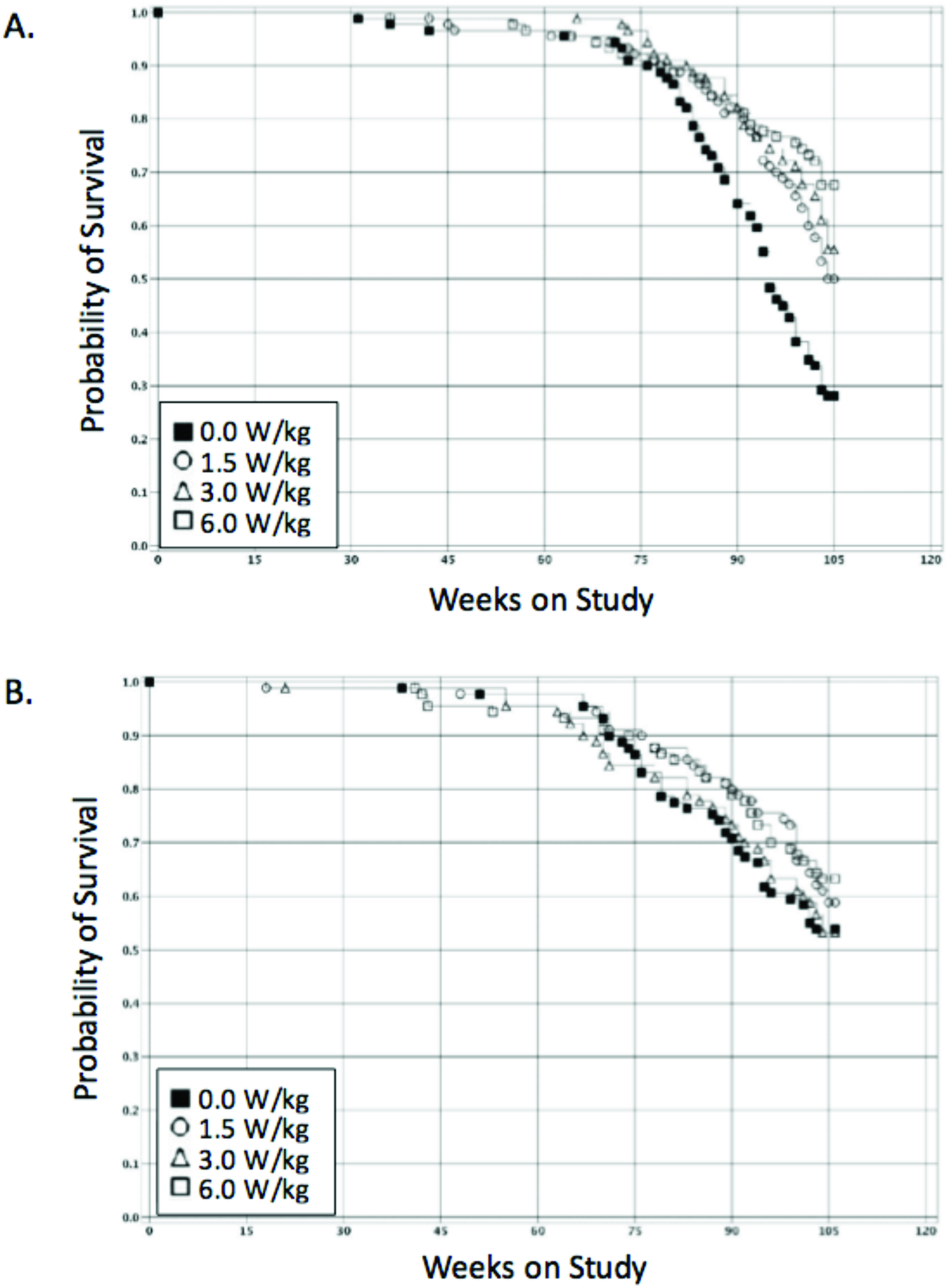
Kaplan-Meier Survival Curves for Male (A) and Female (B) Rats Exposed to Whole Body GSM-Modulated RFR for 2 Years

Survival was also slightly lower in control females than in females exposed to 1.5 or 6 W/kg GSM-modulated RFR. In rats exposed to CDMA-modulated RFR, survival was higher in all groups of exposed males and in the 6 W/kg females compared to controls (Figure 4).

**Figure 4.**
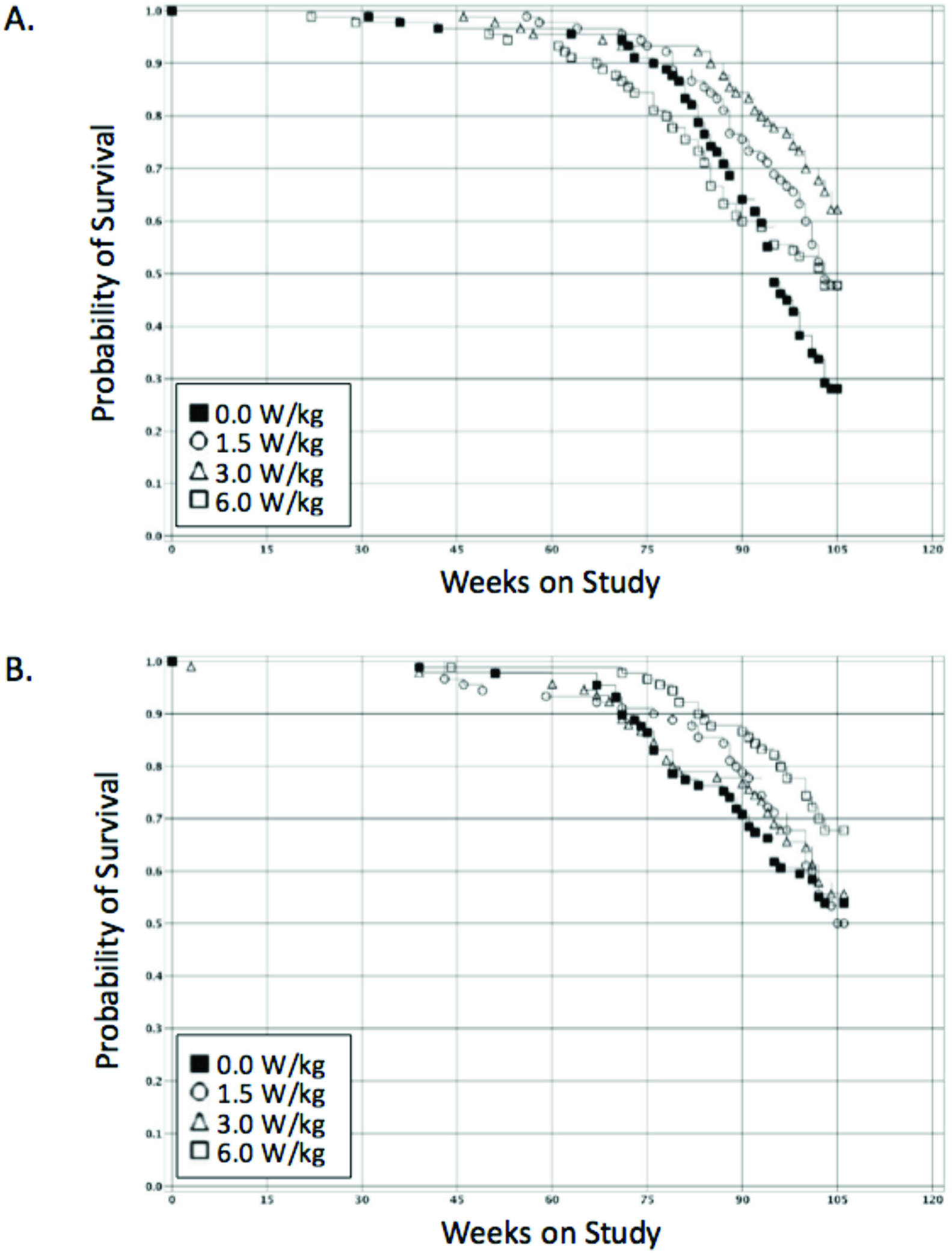
Kaplan-Meier Survival Curves for Male (A) and Female (B) Rats Exposed to Whole Body CDMA-Modulated RFR for 2 Years

### Brain

A low incidence of malignant gliomas and glial cell hyperplasia was observed in all groups of male rats exposed to GSM-modulated RFR (Table 1). In males exposed to CDMA-modulated RFR, a low incidence of malignant gliomas occurred in rats exposed to 6 W/kg (Table 1). Glial cell hyperplasia was also observed in the 1.5 W/kg and 6 W/kg CDMA-modulated exposure groups. No malignant gliomas or glial cell hyperplasias were observed in controls. There was not a statistically significant difference between the incidences of lesions in exposed male rats compared to control males for any of the GSM-or CDMA-modulated RFR groups. However, there was a statistically significant positive trend in the incidence of malignant glioma (p < 0.05) for CDMA-modulated RFR exposures.

**Table 1.** Incidence of brain lesions in male Hsd:Sprague Dawley^®^ SD^®^ (Harlan) rats exposed to GSM-or CDMA-modulated RFR^§^

|  | Control | GSM |  |  | CDMA |  |  |
| --- | --- | --- | --- | --- | --- | --- | --- |
|  | 0<br>W/kg | 1.5<br>W/kg | 3<br>W/kg | 6<br>W/kg | 1.5<br>W/kg | 3<br>W/kg | 6<br>W/kg |
| Number examined | 90 | 90 | 90 | 90 | 90 | 90 | 90 |
| Malignant glioma<sup>†‡</sup> | 0* | 3 (3.3%) | 3 (3.3%) | 2 (2.2%) | 0 | 0 | 3 (3.3%) |
| Glial cell hyperplasia | 0 | 2 (2.2%) | 3 (3.3%) | 1 (1.1%) | 2 (2.2%) | 0 | 2 (2.2%) |

<sup>§</sup> Data presented as number of animals per group with lesions (percentage of animals per group with lesions).
\* Significant SAR-dependent trend for CDMA exposures by poly-6 ( $p < 0.05$ ). See appendix B
<sup>†</sup> Poly-6 survival adjusted rates for malignant gliomas were 0/53.48 in controls; GSM: 3/67.96 (4.4%), 3/72.10 (4.2%), and 2/72.65 (2.8%) in the 1.5, 3, and 6 W/kg groups, respectively; CDMA: 0/65.94, 0/73.08, and 3/57.49 (5.2%) for the 1.5, 3, and 6 W/kg groups, respectively.
<sup>‡</sup> Historical control incidence in NTP studies: 11/550 (2.0%), range 0-8%

In females exposed to GSM-modulated RFR, a malignant glioma was observed in a single rat exposed to 6 W/kg, and glial cell hyperplasia was observed in a single rat exposed to 3 W/kg (Table 2). In females exposed to CDMA-modulated RFR, malignant gliomas were observed in two rats exposed to 1.5 W/kg. Glial cell hyperplasia was observed in one female in each of the CDMA-modulation exposure groups (1.5, 3, and 6 W/kg). There was no glial cell hyperplasia or malignant glioma observed in any of the control females. Detailed descriptions of the malignant gliomas and glial cell hyperplasias are presented in Appendix C.

**Table 2.**
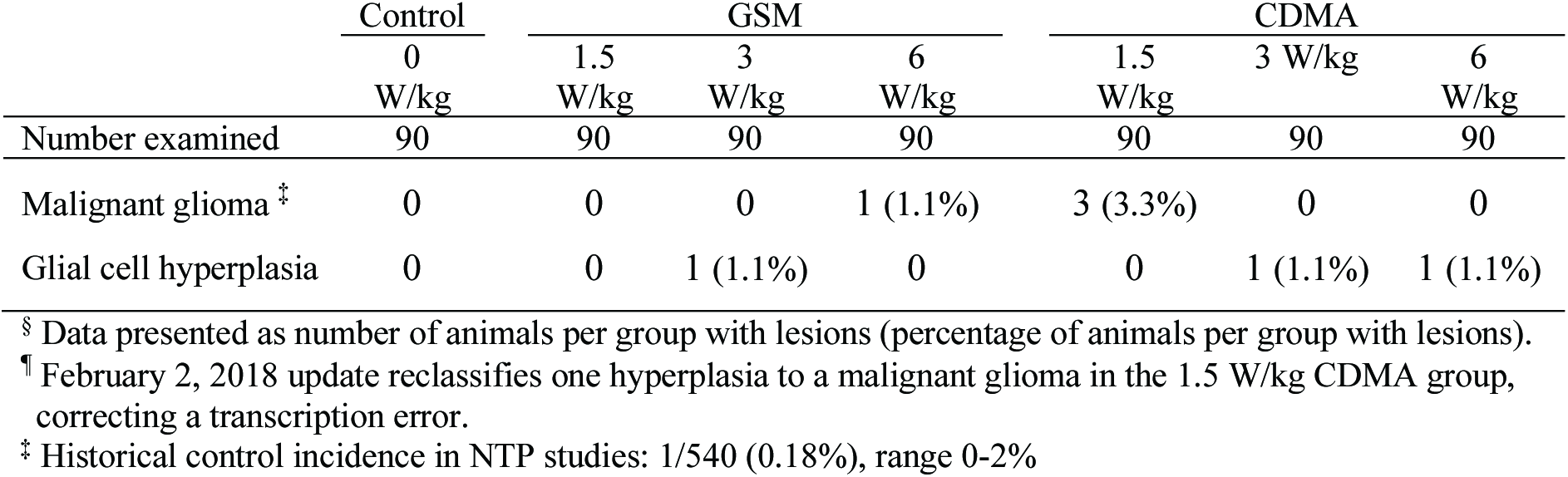
Incidence of brain lesions in female Hsd:Sprague Dawley^®^ SD^®^ (Harlan) rats exposed to GSM-or CDMA-modulated RFR^§,¶^

### Heart

Cardiac schwannomas were observed in male rats in all exposed groups of both GSM-and CDMA-modulated RFR, while none were observed in controls (Table 3). For both modulations (GSM and CDMA), there was a significant positive trend in the incidence of schwannomas of the heart with respect to exposure SAR. Additionally, the incidence of schwannomas in the 6 W/kg males was significantly higher in CDMA-modulated RFR-exposed males compared to controls. The incidence of schwannomas in the 6 W/kg GSM-modulated RFR-exposed males was higher, but not statistically significant (p = 0.052) compared to controls. Schwann cell hyperplasia of the heart was also observed in three males exposed to 6 W/kg CDMA-modulated RFR. In the GSM-modulation exposure groups, a single incidence of Schwann cell hyperplasia was observed in a 1.5 W/kg male and 2 in the 6W/kg group.

**Table 3.** Incidence of heart lesions in male Hsd:Sprague Dawley^®^ SD^®^ (Harlan) rats exposed to GSM-or CDMA-modulated cell phone RFR^§^

|  | Control | GSM |  |  | CDMA |  |  |
| --- | --- | --- | --- | --- | --- | --- | --- |
|  | 0<br>W/kg | 1.5<br>W/kg | 3<br>W/kg | 6<br>W/kg | 1.5<br>W/kg | 3<br>W/kg | 6<br>W/kg |
| Number examined | 90 | 90 | 90 | 90 | 90 | 90 | 90 |
| Schwannoma †‡ | 0* | 2 (2.2%) | 1 (1.1%) | 5 (5.5%) | 2 (2.2%) | 3 (3.3%) | 6 (6.6%)** |
| Schwann cell hyperplasia | 0 | 1 (1.1%) | 0 | 2 (2.2%) | 0 | 0 | 3 (3.3%) |

§ Data presented as number of animals per group with lesions (percentage of animals per group with lesions).
\* Significant SAR level-dependent trend for GSM and CDMA by poly-3 ( $p < 0.05$ ). See appendix B
\*\* Significantly higher than controls by poly-3 ( $p < 0.05$ )
† Poly-3 survival adjusted rates for schwannomas were 0/65.47 in controls; GSM: 2/74.87 (2.7%), 1/77.89 (1.3%), and 5/78.48 (6.4%) in the 1.5, 3, and 6 W/kg groups, respectively; CDMA: 2/74.05 (2.7%), 3/78.67 (3.8%), and 6/67.94 (8.8%) for the 1.5, 3, and 6 W/kg groups, respectively.
‡ Historical control incidence in NTP studies: 9/699 (1.3%) range 0-6%

In females, schwannomas of the heart were also observed at 3 W/kg GSM-modulated RFR and 1.5 and 6 W/kg CDMA-modulated RFR. Schwann cell hyperplasia was observed in one female at 6 W/kg GSM, and in each of the CDMA-modulation exposure groups (1.5, 3, and 6 W/kg).

**Table 4.** Incidence of heart lesions in female Hsd:Sprague Dawley^®^ SD^®^ (Harlan) rats exposed to GSM-or CDMA-modulated cell phone RFR^§^

|  | Control | GSM |  |  | CDMA |  |  |
| --- | --- | --- | --- | --- | --- | --- | --- |
|  | 0<br>W/kg | 1.5 W/kg | 3<br>W/kg | 6 W/kg | 1.5<br>W/kg | 3<br>W/kg | 6<br>W/kg |
| Number examined | 90 | 90 | 90 | 90 | 90 | 90 | 90 |
| Schwannoma† | 0 | 0 | 2 (2.2%) | 0 | 2 (2.2%) | 0 | 2 (2.2%) |
| Schwann cell hyperplasia | 0 | 0 | 0 | 1 (1.1%) | 1 (1.1%) | 1 (1.1%) | 1 (1.1%) |

§ Data presented as number of animals per group with tumors (percentage of animals per group with tumors).
† Historical control incidence in NTP studies: 4/699 (0.6 %), range 0-4%

Schwann cells are present in the peripheral nervous system and are distributed throughout the whole body, not just in the heart. Therefore, organs other than the heart were examined for schwannomas and Schwann cell hyperplasia. Several occurrences of schwannomas were observed in the head, neck, and other sites throughout the body of control and GSM and CDMA RFR-exposed male rats. In contrast to the significant increase in the incidence of schwannomas in the heart of exposed males, the incidence of schwannomas observed in other tissue sites of exposed males (GSM and CDMA modulations) was not significantly different than in controls (Table 5). Additionally, Schwann cell hyperplasia was not observed in any tissues other than the heart. The combined incidence of schwannomas from all sites was generally higher in GSM-and CDMA-modulated RFR exposed males, but not significantly different than in controls. The Schwann cell response to RFR appears to be specific to the heart of male rats.

**Table 5.** Incidence of schwannomas in male Hsd:Sprague Dawley^®^ SD^®^ (Harlan) rats exposed to GSM-or CDMA-modulated RFR^§,¶^

|  | Control | GSM |  |  | CDMA |  |  |
| --- | --- | --- | --- | --- | --- | --- | --- |
|  | 0 | 1.5 | 3 | 6 | 1.5 | 3 | 6 |
|  | W/kg | W/kg | W/kg | W/kg | W/kg | W/kg | W/kg |
| Number examined | 90 | 90 | 90 | 90 | 90 | 90 | 90 |
| Heart<sup>‡</sup> | 0* | 2 (2.2%) | 1 (1.1%) | 5 (5.5%) | 2 (2.2%) | 3 (3.3%) | 6 (6.6%)** |
| Other sites<sup>†</sup> | 3 (3.3%) | 1 (1.1%) | 4 (4.4%) | 2 (2.2%) | 2 (2.2%) | 1 (1.1%) | 2 (2.2%) |
| All sites (total) | 3 (3.3%) | 3 (3.3%) | 5 (5.5%) | 7 (7.7%) | 4 (4.4%) | 4 (4.4%) | 8 (8.8%) |

<sup>§</sup> Data presented as number of animals per group with tumors (percentage of animals per group with tumors).
<sup>¶</sup> February 2, 2018 update adds one additional tumor in the 6 W/kg CDMA group, identified as a result of additional pathology reviews of other sites.
\* Significant SAR level-dependent trend for GSM and CDMA, poly 3 test ( $p < 0.05$ )
\*\* Significantly higher than controls, poly-3 test ( $p < 0.05$ )
<sup>‡</sup> Historical control incidence in NTP studies: 9/699 (1.3%), range 0-6%
<sup>†</sup> Mediastinum, thymus, and fat

In female rats, there was no statistically significant or apparent exposure-related effect on the incidence of schwannomas in the heart or the combined incidence in the heart or other sites (Table 6).

**Table 6.** Incidence of schwannomas in female Hsd:Sprague Dawley^®^ SD^®^ (Harlan) rats exposed to GSM-or CDMA-modulated RFR^§^

| Schwannoma site | Control | GSM |  |  |  | CDMA |  |  |
| --- | --- | --- | --- | --- | --- | --- | --- | --- |
|  | 0<br>W/kg | 1.5<br>W/kg | 3<br>W/kg | 6<br>W/kg |  | 1.5<br>W/kg | 3<br>W/kg | 6<br>W/kg |
| Number examined | 90 | 90 | 90 | 90 |  | 90 | 90 | 90 |
| Heart<sup>‡</sup> | 0 | 0 | 2 (2.2%) | 0 |  | 2 (2.2%) | 0 | 2 (2.2%) |
| Other sites<sup>†</sup> | 4 (4.4%) | 1 (1.1%) | 3 (3.3%) | 2 (2.2%) |  | 0 | 2 (2.2%) | 2 (2.2%) |
| All sites (total) | 4 (4.4%) | 1 (1.1%) | 5 (5.5%) | 2 (2.2%) |  | 2 (2.2%) | 2 (2.2%) | 4 (4.4%) |

§ Data presented as number of animals per group with tumors (percentage of animals per group with tumors).
‡ Historical control incidence in NTP studies: 4/699 (0.6%), range 0-4%
† Ovary, uterus, vagina, thymus, abdomen, and clitoral gland

## DISCUSSION

The two tumor types, which are the focus of this report, are malignant gliomas of the brain and schwannomas of the heart. Glial cells are a collection of specialized, non-neuronal, support cells whose functions include maintenance of homeostasis, formation of myelin, and providing support and protection for neurons of the peripheral nervous system (PNS) and the central nervous system (CNS). In the CNS, glial cells include astrocytes, oligodendrogliocytes, microglial cells, and ependymal cells. Schwann cells are classified as glial cells of the PNS. In the PNS, Schwann cells produce myelin and are analogous to oligodendrocytes of the CNS. Generally, glial neoplasms in the rat are aggressive, poorly differentiated, and usually classified as malignant.

In the heart, exposure to GSM or CDMA modulations of RFR in male rats resulted in a statistically significant, positive trend in the incidence of schwannomas. There was also a statistically significant, pairwise increase at the highest CDMA exposure level tested compared to controls. Schwann cell hyperplasias also occurred at the highest exposure level of CDMA-modulated RFR. The intracardiac schwannomas in male rats were not observed in animals from the same litter. Schwann cell hyperplasia in the heart may progress to cardiac schwannomas. No Schwann cell hyperplasias or schwannomas of the heart were observed in the single, common control group of male rats. The historical control rate of schwannomas of the heart in male Harlan Sprague Dawley rats is 1.30% (7/539) and ranges from 0-6% for individual NTP studies (Table D2, Appendix D). The 5.5-6.6% observed in the 6 W/kg GSM-and CDMA-modulated RFR groups exceeds the historical incidence, and approaches or exceeds the highest rate observed in a single study (6%). The increase in the incidence of schwannomas in the heart of male rats in this study is likely the result of whole-body exposures to GSM-or CDMA-modulated RFR.

In the brain, there was a significant, positive trend in the incidences of malignant gliomas in males exposed to CDMA-modulated RFR, and a low incidence was observed in males at all exposure levels of GSM-modulated RFR that was not statistically different than in control males. The male rats in which gliomas were observed were not from the same litter. Glial cell hyperplasia, a preneoplastic lesion distinctly different from gliosis, was also observed at low incidences in rats exposed to either GSM or CDMA modulation. Glial cell hyperplasia may progress to malignant glioma. Neither of these lesions was observed in the control group of male rats. Although not observed in the current control group, malignant gliomas have been observed in control male Harlan Sprague Dawley rats from other completed NTP studies. Currently in males, the historical control rate of malignant glioma for those studies is 2.0% (11/550) and ranges from 0-8% for individual studies (Table D1, Appendix D). The 2.2-3.3% observed in all of the GSM-modulation groups and in the 6 W/kg CDMA-modulated group only slightly exceeds the mean historical control rate and falls within the observed range.

The survival of the control group of male rats in the current study (28%) was relatively low compared to other recent NTP studies in Hsd:Sprague Dawley^®^ SD^®^ (Harlan) rats (average 47%, range 24-72%). If malignant gliomas or schwannomas are late-developing tumors, the absence of these lesions in control males in the current study could conceivably be related to the shorter longevity of control rats in this study. Appendix E lists the time on study for each animal with a malignant glioma or heart schwannoma. Most of the gliomas were observed in animals that died late in the study, or at the terminal sacrifice. However, a relatively high number of the heart schwannomas in exposed groups were observed by 90 weeks into the study, a time when approximately 60 of the 90 control male rats remained alive and at risk for developing a tumor.

## CONCLUSIONS

Under the conditions of these 2-year studies, the hyperplastic lesions and glial cell neoplasms of the heart and brain observed in male rats are considered likely the result of whole-body exposures to GSM-or CDMA-modulated RFR. There is higher confidence in the association between RFR exposure and the neoplastic lesions in the heart than in the brain. No biologically significant effects were observed in the brain or heart of female rats regardless of modulation.

## NEXT STEPS

The results reported here are limited to select findings of concern in the brain and heart and do not represent a complete reporting of all findings from these studies of cell phone RFR. The complete results for all NTP studies on the toxicity and carcinogenicity of GSM and CDMA-modulated RFR are currently being reviewed and evaluated according to the established NTP process and will be reported together with the current findings in two forthcoming NTP Technical Reports. Given the large scale and scope of these studies, completion of this process is anticipated by fall 2017, and the draft NTP Technical Reports are expected to be available for peer review and public comment in early 2018. We anticipate that the results from a series of initial studies investigating the tolerance to various power levels of RFR, including measurements of body temperatures in both sexes of young and old rats and mice and in pregnant female rats, will be published in the peer-reviewed literature in early 2018 as well.

## APPENDIX A – CONTRIBUTORS NTP CONTRIBUTORS

Participated in the evaluation and interpretation of results and the reporting of findings.

M.E. Wyde, Ph.D. (NTP study scientist)

M.F. Cesta, D.V.M., Ph.D. (NTP pathologist)

C.R. Blystone, Ph.D.

J.R. Bucher, Ph.D.

S.A. Elmore, D.V.M., M.S.

P.M. Foster, Ph.D.

M.J. Hooth, Ph.D.

G.E. Kissling, Ph.D.

D.E. Malarkey, D.V.M., Ph.D.

R.C. Sills, D.V.M., Ph.D.

M.D. Stout, Ph.D.

N.J. Walker, Ph.D.

K.L. Witt, M.S.

M.S. Wolfe, Ph.D.

## APPENDIX B STATISTICAL ANALYSIS

## Appendix B1 Statistical Methods

The Poly-k test (Bailer and Portier, 1988; Portier and Bailer, 1989; Piegorsch and Bailer, 1997) was used to assess neoplasm prevalence. This test is a survival-adjusted quantal-response procedure that modifies the Cochran-Armitage linear trend test to take survival differences into account. More specifically, this method modifies the denominator in the quantal estimate of lesion incidence to approximate more closely the total number of animal years at risk. For analysis of lesion incidence at a given site, each animal is assigned a risk weight. This value is one if the animal had a lesion at that site or if it survived until terminal sacrifice; if the animal died prior to terminal sacrifice and did not have a lesion at that site, its risk weight is the fraction of the entire study time that it survived, raised to the kth power. This method yields a lesion prevalence rate that depends only upon the choice of a shape parameter, k, for a Weibull hazard function describing cumulative lesion incidence over time (Bailer and Portier, 1988). A further advantage of the Poly-k method is that it does not require lesion lethality assumptions.

Unless otherwise specified, the NTP uses a value of k=3 in the analysis of site-specific lesions (Portier et al., 1986). Bailer and Portier (1988) showed that the Poly-3 test gives valid results if the true value of k is anywhere in the range from 1 to 5. In addition, Portier et al. (1986) modeled a collection of relatively common tumors observed in control animals from two-year NTP rodent carcinogenicity studies, showing that the Weibull distribution with values of k ranging between 1 and 5 was a reasonable fit to tumor incidence in most cases. In cases of early tumor onset or late tumor onset, however, k=3 may not be the optimal choice. Tumors with early onset would require a value of k much less than 3, while tumors with late onset would require a value of k much greater than 3. In the current studies, malignant brain gliomas occurred only in animals surviving more than 88% of the length of the study. For these brain tumors, a Weibull distribution with k=6 is a better fit to survival time than with k=3 (Portier, 1986). Malignant schwannomas of the heart occurred in animals surviving at least 65% of the length of the study; a Weibull distribution with k=3 adequately fits these heart tumor incidences. Therefore, poly-6 tests were used for analyses of brain tumors and poly-3 tests were used for schwannomas. Variation introduced by the use of risk weights, which reflect differential mortality, was accommodated by adjusting the variance of the Poly-k statistic as recommended by Bieler and Williams (1993) and a continuity correction modified from Thomas et al. (1977) was applied.

Tests of significance for tumors and nonneoplastic lesions included pairwise comparisons of each dosed group with controls and a test for an overall dose-related trend. Continuity-corrected Poly-k tests were used in the analysis of lesion incidence, and reported P values are one sided.

Body weights and litter weights were compared to the control group using analysis of variance and Dunnett’s test (1955). The probability of survival was estimated by the product-limit procedure of Kaplan and Meier (1958). Statistical analyses for possible exposure-related effects on survival used Cox’s (1972) method for testing two groups for equality and Tarone’s (1975) life table test to identify exposure-related trends. Survival analysis p-values are two-sided.

## Appendix B2 Statistical Significance of Rare Tumors in a Low Powered Situation - NTP View

In Dr. Lauer’s review (Appendix G1, p 47) he states: “The low power implies that there is a high risk of false positive findings, especially since the epidemiological literature questions the purported association between cell phone exposure and cancer.” He kindly provided three additional references (cited below) to justify his statement.

The three papers make the argument that in studies that have low power to detect an effect, a significant finding (*p* value <0.05) is more likely to be a false positive than a true positive. In some cases this may be correct, but for rare tumors, as observed in the current study, it is very unlikely that the significant findings are false positives. One reason for this is that the actual significance level of the tests is not 0.05; it is <u>much</u> less, as illustrated below. Another reason is the introduction and use of rates of true prevalence of effects, which we consider first.

A few definitions are useful:

The significance level of the test, *α*, is the probability of a false positive. The power of the test, *1 – β*, is the probability of a true positive. Following the notation in the Button *et al.* (2013) reference, the Positive Predictive Value, *PPV*, is the probability that a statistically significant result is a true positive.

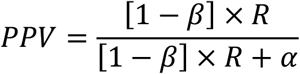

This expression involves *R*, which is the pre-study odds that the tested effect is a true effect. That is,

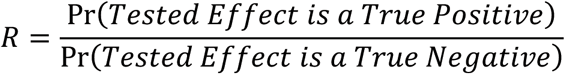

As illustrated below in the statistical examples, *R* is a major modifying factor governing whether a result with a significant *p* value is appropriately considered a true or false positive. The selection of an appropriate *R*, or expected odds of an effect, in this case a carcinogenic effect*, permits the introduction of bias in the interpretation of the *p* values in this report on the NTP cancer studies of radiofrequency radiation.

For example, *R* could be pre-assigned as the expected odds of a positive cancer finding at any site in male or female rats or mice (as in scenario 1 below where R = 1.2), or as the odds of a positive cancer finding in only the male rat (as in scenario 2 below where R = 0.54). In these two cases, the chances of our findings being <u>true</u> positives (*PPV*) are very high (>94%), despite the low power of the current study to detect such an effect.

*R* could alternatively be pre-assigned as the odds of seeing these specific tumor types occur only in the brain and/or heart of male rats. In this case, because gliomas and schwannomas have only been part of two positive calls for cancer in male rats in any of the prior studies of <u>chemical</u> agents that NTP has performed, *R* is much lower (0.035) and the chance of a <u>false</u> positive finding is indeed high. But, this is the case even if the power to detect an effect is high; i.e., the conclusion of false positivity is driven more by the *a priori* expectation of a low odds of occurrence of an effect than it is by the power to detect the effect. At high values of *R* (*R*>1), all outcomes that are p<0.05, regardless of the actual power, will generally be considered “true positives.” At very low values of *R* (*R*<0.01), even an outcome from a study that has high power, will be considered a false positive (*PPV*<0.2).

Dr. Lauer’s comment, “the epidemiological literature questions the purported association between cell phone exposure and cancer,” would place the expected *R* close to zero. As can be seen at very low values of *R* approaching zero, all findings, regardless of the power to detect them, will be considered false positives. On the other hand, if one is open to the possibility that *R* is in fact non-zero, then the findings need to become part of the public discussion over the safety of exposures to RFR.

**STATISTICAL EXAMPLES**

To illustrate, suppose that the background tumor rate is 1.5%, which is similar to the rate of schwannomas in the hearts of male rats in the NTP historical control database (1.3%), and that there are two groups: Control and Treated, with n = 90 animals per group. Further suppose that the null hypothesis that tumor rates are the same in the two groups, H0, is tested against the alternative hypothesis that the Treated group has a higher rate, Ha, using a one-sided Fisher’s exact test. We reject the null hypothesis if *p* is less than 0.05. The <u>actual</u> significance level of this test, *α*, is the probability of rejecting the null hypothesis when is it actually true. In other words,

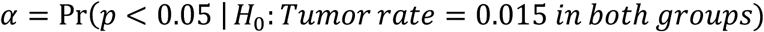

By Fisher’s exact test, *p* < 0.05 if there are

- 0 tumors in the Control group and 5 or more in the Treated group, or
- 1 tumor in the Control group and 7 or more in the Treated group, or
- 2 tumors in the Control group and 8 or more in the Treated group, or
- 3 tumors in the Control group and 10 or more in the Treated group, or
- ….

The probability of making a Type I error (false positive decision, rejecting H0 when it is true) is:

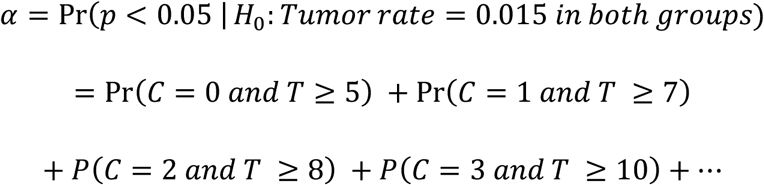

Using binomial probabilities,

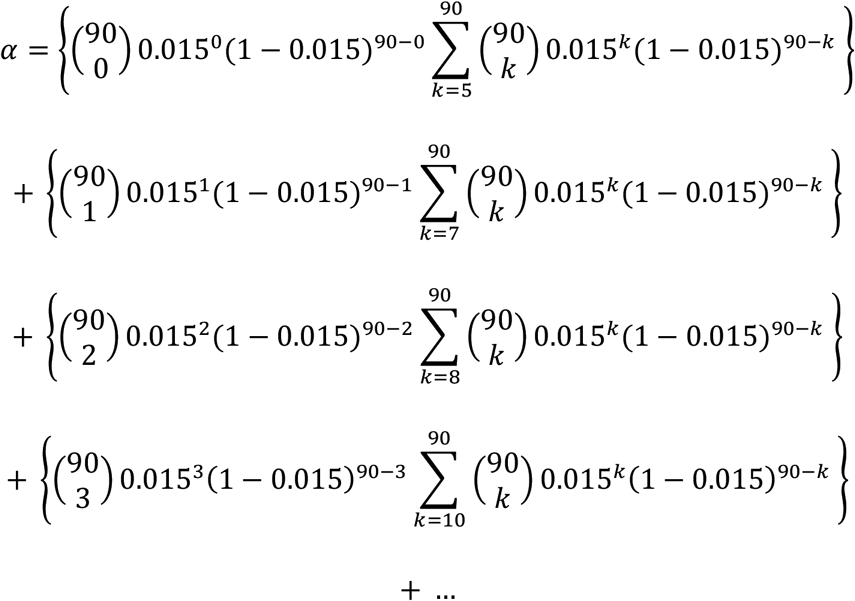

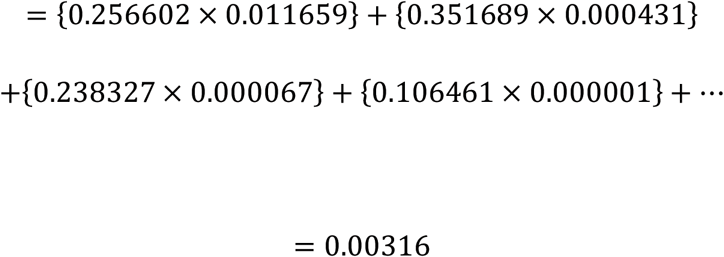

Thus, when the tumor rate is low and the decision rule is to reject H0 when the one-sided Fisher’s exact p-value is less than 0.05, the actual false positive rate, α, is 0.0032.

This significance level can be used to calculate the probability that a significant result is a true positive, Positive Predictive Value (*PPV*). Following the Button *et al.* (2013) paper’s notation:

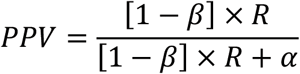

Suppose that the power, *1 − β*, is 10%. *R* as defined in Button *et al.* is the pre-study odds of a true positive. There are several possible ways to estimate *R*:

1. Among the 595 NTP studies that have a determination about carcinogenesis, 326 concluded that the test article was carcinogenic^*^. The pre-study odds of a carcinogenic effect, *R*, is 326/(595-326) = 1.2119. Thus, the probability that a significant test represents a true positive is

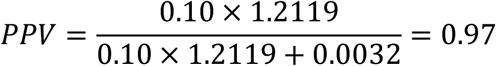 This says that, under the low power/low tumor rate conditions described above, if a test is significant at the 0.05 level, it almost certainly indicates a real carcinogenic effect.
2. Alternatively, among the 580 NTP studies that involved male rats, 203 concluded that the test article was carcinogenic in male rats; thus, *R* = 203/(580 - 203) = 0.538 and *PPV* = 0.94.
3. If there is no prior information and it is thought that it is as equally likely that there is a real effect as it is that there is no effect, then *R* = 1 and *PPV* = 0.97.

Furthermore, the relationship between *R* and *PPV* can be rearranged to solve for *R*,

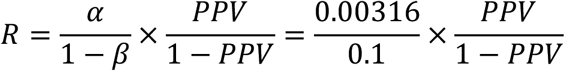

In this low power/low tumor rate situation, *R* could be as low as 0.28 and the *PPV* would be at least 90%, or *R* could be as low as 0.13 and the *PPV* would be at least 80%.

Dr. Grace Kissling, NTP study statistician, provided the statistical illustrations. Also Dr. Shyamal Peddada, Acting Chief, Biostatistics and Computational Biology Branch, NIEHS, has reviewed and concurs with her interpretation of the issues posited by the papers and the explanation of why they do not diminish the importance of the findings from the current study.

We have not assigned a specific level of evidence to the NTP RFR study, as it is not complete. Rather, we evaluated the partial study findings and concluded that the tumors highlighted are “likely” related to the RFR exposure.

**EXPLANATION OF LEVELS OF EVIDENCE OF CARCINOGENIC ACTIVITY**

The National Toxicology Program describes the results of individual experiments on a chemical agent and notes the strength of the evidence for conclusions regarding each study. Negative results, in which the study animals do not have a greater incidence of neoplasia than control animals, do not necessarily mean that a chemical is not a carcinogen, inasmuch as the experiments are conducted under a limited set of conditions. Positive results demonstrate that a chemical is carcinogenic for laboratory animals under the conditions of the study and indicate that exposure to the chemical has the potential for hazard to humans. Other organizations, such as the International Agency for Research on Cancer, assign a strength of evidence for conclusions based on an examination of all available evidence, including animal studies such as those conducted by the NTP, epidemiologic studies, and estimates of exposure. Thus, the actual determination of risk to humans from chemicals found to be carcinogenic in laboratory animals requires a wider analysis that extends beyond the purview of these studies.

Five categories of evidence of carcinogenic activity are used in the Technical Report series to summarize the strength of evidence observed in each experiment: two categories for positive results **(clear evidence and some evidence)**; one category for uncertain findings **(equivocal evidence)**; one category for no observable effects **(no evidence)**; and one category for experiments that cannot be evaluated because of major flaws **(inadequate study)**. These categories of interpretative conclusions were first adopted in June 1983 and then revised on March 1986 for use in the Technical Report series to incorporate more specifically the concept of actual weight of evidence of carcinogenic activity. For each separate experiment (male rats, female rats, male mice, female mice), one of the following five categories is selected to describe the findings. These categories refer to the strength of the experimental evidence and not to potency or mechanism.

- **Clear evidence** of carcinogenic activity is demonstrated by studies that are interpreted as showing a dose-related (i) increase of malignant neoplasms, (ii) increase of a combination of malignant and benign neoplasms, or (iii) marked increase of benign neoplasms if there is an indication from this or other studies of the ability of such tumors to progress to malignancy.
- **Some evidence** of carcinogenic activity is demonstrated by studies that are interpreted as showing a chemical-related increased incidence of neoplasms (malignant, benign, or combined) in which the strength of the response is less than that required for clear evidence.
- **Equivocal evidence** of carcinogenic activity is demonstrated by studies that are interpreted as showing a marginal increase of neoplasms that may be chemical related.
- **No evidence** of carcinogenic activity is demonstrated by studies that are interpreted as showing no chemical-related increases in malignant or benign neoplasms
- **Inadequate study** of carcinogenic activity is demonstrated by studies that, because of major qualitative or quantitative limitations, cannot be interpreted as valid for showing either the presence or absence of carcinogenic activity. For studies showing multiple chemical-related neoplastic effects that if considered individually would be assigned to different levels of evidence categories, the following convention has been adopted to convey completely the study results. In a study with clear evidence of carcinogenic activity at some tissue sites, other responses that alone might be deemed some evidence are indicated as “were also related” to chemical exposure. In studies with clear or some evidence of carcinogenic activity, other responses that alone might be termed equivocal evidence are indicated as “may have been” related to chemical exposure. When a conclusion statement for a particular experiment is selected, consideration must be given to key factors that would extend the actual boundary of an individual category of evidence. Such consideration should allow for incorporation of scientific experience and current understanding of long-term carcinogenesis studies in laboratory animals, especially for those evaluations that may be on the borderline between two adjacent levels. These considerations should include:

- adequacy of the experimental design and conduct;
- occurrence of common versus uncommon neoplasia;
- progression (or lack thereof) from benign to malignant neoplasia as well as from preneoplastic to neoplastic lesions;
- some benign neoplasms have the capacity to regress but others (of the same morphologic type) progress. At present, it is impossible to identify the difference. Therefore, where progression is known to be a possibility, the most prudent course is to assume that benign neoplasms of those types have the potential to become malignant;
- combining benign and malignant tumor incidence known or thought to represent stages of progression in the same organ or tissue;
- latency in tumor induction;
- multiplicity in site-specific neoplasia;
- metastases;
- supporting information from proliferative lesions (hyperplasia) in the same site of neoplasia or other experiments (same lesion in another sex or species);
- presence or absence of dose relationships;
- statistical significance of the observed tumor increase;
- concurrent control tumor incidence as well as the historical control rate and variability for a specific neoplasm;
- survival-adjusted analyses and false positive or false negative concerns;
- structure-activity correlations; and
- in some cases, genetic toxicology.

## APPENDIX C PATHOLOGY

Pathology data presented in this report on cell phone RFR were subjected to a rigorous peer review process. The primary goal of the NTP peer-review process is to reach consensus agreement on treatment-related findings, confirm the diagnosis of all neoplasms, and confirm any unusual lesions. At study termination, a complete necropsy and histopathology evaluation was conducted on every animal. The initial pathology examination was performed by a veterinary pathologist, who recorded all neoplastic and nonneoplastic lesions. This examination identified several potential treatment-related lesions in target organs of concern (brain and heart), which were chosen for immediate review.^1^ The initial findings of glial cell tumors and hyperplasias in the brain and schwannomas, Schwann call hyperplasia, and schwannomas from all sites were subjected to an expedited, multilevel NTP pathology peer-review process. The data were locked^2^ prior to receipt of the finalized, study-laboratory reports to ensure that the raw data did not change during the review.

The pathology peer review consisted of a quality assessment (QA) review of all slides with tissues from the central nervous system (7 sections of brain and 3 sections of spinal cord), trigeminal nerve and ganglion, and heart. Additionally, the schwannomas of the head and neck region were reviewed. The QA review of the central nervous system and head and neck schwannomas was performed by Dr. Margarita Gruebbel of Experimental Pathology Laboratories, Inc. (EPL), and the QA review of the hearts and trigeminal nerves and ganglia was performed by Dr. Cynthia Shackelford, EPL.

The QA review pathologists then met with Dr. Mark Cesta, NTP pathologist for these studies, and Dr. David Malarkey, head of the NTP Pathology Group, to review lesions and select slides for the Pathology Working Group (PWG) reviews. All PWG reviews were conducted blinded with respect to treatment group and only identified the test articles as “test agent A” or “test agent B”. Due to the large number of slides for review, the PWG was held in three separate sessions:

- January 29, 2016, for review of glial lesions in the brain and Schwann cell lesions in the heart
- February 11, 2016, for review of schwannomas of the head and neck
- February 12, 2016, for review of granular cell lesions of the brain

The reviewing PWG pathologists largely agreed on the diagnostic criteria for the lesions and on the diagnoses of schwannomas in the head and neck, and granular cell lesions in the brain. However, there was much discussion on the criteria for differentiating glial cell hyperplasia from malignant glioma and Schwann cell hyperplasia from schwannoma. The lack of PWG agreement on definitive criteria for the glial cell and Schwann cell lesions, and the requirement for a high level of confidence in the diagnoses prompted NTP to convene two additional PWGs (organized and conducted by the NTP pathologist, Dr. Mark Cesta) with selected experts in the organ under review. These second level PWG reviews were also conducted as noted above and held in two separate sessions:

- February 25, 2016, for review of glial lesions in the brain
- March 3, 2016, for review of cardiac schwannomas, schwannomas in other organs (except the head and neck), and right ventricular degeneration

In both PWGs, the participants came to consensus on the diagnoses of the lesions and the criteria used for those diagnoses. Participants of the individual PWGs are listed below.

**Table C-1.** NTP Pathology Working Group (PWG) Attendees

| PWG member | Affiliation |
| --- | --- |
| *January 29, 2016 - Evaluated glial lesions in the brain and Schwann cell lesions in the heart* |  |
| A.E. Brix, D.V.M., Ph.D. | Experimental Pathology Laboratories, Inc. RTP, NC |
| M.F. Cesta, D.V.M., Ph.D. | National Institute of Environmental Health Sciences<br>(NTP study pathologist) |
| S.A. Elmore, D.V.M., MS | National Institute of Environmental Health Sciences |
| G.P. Flake, M.D. | National Institute of Environmental Health Sciences |
| R.H. Garman, D.V.M. | Consultants in Veterinary Pathology, Inc. Monroeville, PA |
| M.M. Gruebbel, D.V.M., Ph.D. | Experimental Pathology Laboratories, Inc. RTP, NC (observer) |
| R.A. Herbert, D.V.M., Ph.D. | National Institute of Environmental Health Sciences |
| J.S. Hoane, D.V.M. | Charles River Laboratories, Inc. Durham, NC (contract study pathologist) |
| K.S. Janardhan, BVSc, MVSc, Ph.D. | Integrated Laboratory System |
| R. Kovi, BVSc, MVSc, Ph.D. | Experimental Pathology Laboratories, Inc. RTP, NC (observer) |
| D.E. Malarkey, D.V.M., Ph.D. | National Institute of Environmental Health Sciences |
| R.A. Miller, D.V.M., Ph.D. | Experimental Pathology Laboratories, Inc. RTP, NC |
| J.P. Morrison, D.V.M. | Charles River Laboratories, Inc. Durham, NC |
| A.R. Pandiri, BVSc & AH, Ph.D. | National Institute of Environmental Health Sciences |
| C.C. Shackelford, D.V.M., Ph.D. | Experimental Pathology Laboratories, Inc. RTP, NC (observer) |
| J.A. Swenberg, D.V.M., Ph.D. | University of North Carolina – Chapel Hill, NC |
| G. Willson, BVMS, Dip RC Path, FRC Path, MRCVS | Experimental Pathology Laboratories, Inc. RTP, NC (PWG coordinator) |
| *February 11, 2016 - Evaluated schwannomas of the head and neck* |  |
| A.E. Brix, D.V.M., Ph.D. | Experimental Pathology Laboratories, Inc. RTP, NC |
| M.F. Cesta, D.V.M., Ph.D. | National Institute of Environmental Health Sciences (NTP study pathologist) |
| S.A. Elmore, D.V.M., MS | National Institute of Environmental Health Sciences |
| G.P. Flake, M.D. | National Institute of Environmental Health Sciences |
| M.M. Gruebbel, D.V.M., Ph.D., | Experimental Pathology Laboratories, Inc. RTP, NC (PWG coordinator) |
| K.S. Janardhan, BVSc, MVSc, Ph.D. | Integrated Laboratory System RTP, NC |
| D.E. Malarkey, D.V.M., Ph.D. | National Institute of Environmental Health Sciences |
| A.R. Pandiri, BVSc & AH, Ph.D. | National Institute of Environmental Health Sciences |
| R.R. Maronpot, D.V.M. | Experimental Pathology Laboratories, Inc. RTP, NC |
| *February 12, 2016 - Evaluated granular cell lesions of the brain* |  |
| A.E. Brix, D.V.M., Ph.D. | Experimental Pathology Laboratories, Inc. RTP, NC |
| M.F. Cesta, D.V.M., Ph.D. | National Institute of Environmental Health Sciences (NTP study pathologist) |
| S.A. Elmore, D.V.M., MS | National Institute of Environmental Health Sciences |
| M.M. Gruebbel, D.V.M., Ph.D., | Experimental Pathology Laboratories, Inc. RTP, NC (PWG coordinator) |
| J.S. Hoane, D.V.M. | Charles River Laboratories, Inc. Durham, NC (contract study pathologist) |
| K.S. Janardhan, BVSc, MVSc, Ph.D. | Integrated Laboratory System RTP, NC |
| A.R. Pandiri, BVSc. & AH, Ph.D. | National Institute of Environmental Health Sciences |
| R.R. Moore, D.V.M. | Integrated Laboratory System RTP, NC |
| *February 25, 2016 - Evaluated glial lesions in the brain* |  |
| D. Bigner, M.D., Ph.D. | Duke University Durham, NC |
| B. Bolon, D.V.M., MS, Ph.D. | GEMpath, Inc. Longmont, CO |
| V. Chen, D.V.M., Ph.D. | National Institute of Environmental Health Sciences (observer) |
| M.F. Cesta, D.V.M., Ph.D. | National Institute of Environmental Health Sciences (PWG coordinator, NTP study pathologist) |
| S.A. Elmore, D.V.M., MS | National Institute of Environmental Health Sciences (observer) |
| G.P. Flake, M.D. | National Institute of Environmental Health Sciences (observer) |
| J.S. Hardisty, D.V.M. | Experimental Pathology Laboratories, Inc. RTP, NC |
| R.A. Herbert, D.V.M., Ph.D., | National Institute of Environmental Health Sciences (observer) |
| R. Kovi, BVSc, MVSc, Ph.D. | Experimental Pathology Laboratories, Inc. (observer) |
| P.B. Little, D.V.M. | Experimental Pathology Laboratories, Inc. |
| D.E. Malarkey, D.V.M., Ph.D. | National Institute of Environmental Health Sciences |
| J.P. Morrison, D.V.M., Ph.D. | Charles River Laboratories, Inc. |
| A. Sharma, BVSc, MVSc, MS, Ph.D. | Covance |
| *March 3, 2016 - Evaluated heart lesions, and schwannomas in other organs (except head and neck)* |  |
| B. Berridge, D.V.M., Ph.D. | GlaxoSmithKline RTP, NC |
| M.C. Boyle, D.V.M., Ph.D. | Amgen Thousand Oaks, CA |
| V. Chen, D.V.M., Ph.D. | National Institute of Environmental Health Sciences (observer) |
| M.F. Cesta, D.V.M., Ph.D. | National Institute of Environmental Health Sciences (PWG coordinator, NTP study pathologist) |
| S.A. Elmore, D.V.M., MS | National Institute of Environmental Health Sciences (observer) |
| M. Elwell, D.V.M., Ph.D. | Covance Chantilly, VA |
| J.R. Hailey, D.V.M. | Covance Chantilly, VA |
| M. Novilla, D.V.M., MS, Ph.D. | SNBL Everett, WA |

**LESION DESCRIPTIONS**

*Brain*

Malignant gliomas were infiltrative lesions, usually of modest size, with indistinct tumor margins. The neoplastic cells were typically very densely packed with more cells than neuropil. The cells were typically small and had round to oval, hyperchromatic nuclei. Mitoses were infrequent. In some of the neoplasms, invasion of the meninges, areas of necrosis surrounded by palisading neoplastic cells, cuffing of blood vessels, and neuronal satellitosis were observed. The malignant gliomas did not appear to arise from any specific anatomic subsite of the brain.

Glial cell hyperplasia consisted of small, proliferative, and poorly demarcated foci of poorly differentiated glial cells that accumulated and invaded into the surrounding parenchyma. In some cases, there was a small amount of perivascular cuffing. The hyperplastic cells appeared morphologically identical to those in the gliomas but were typically less dense with more neuropil than glial cells. There were no necrotic or degenerative elements present, so there was no evidence that the increased number of glial cells was a reaction to brain injury.

*Heart*

The intracardiac schwannomas were either endocardial or myocardial (intramural). The endocardial schwannomas lined the ventricles and atria and invaded into the myocardium. Two morphologic cell types were observed, but indistinct cell margins and eosinophilic cytoplasm were common to both types. Groups of cells with widely spaced small, round nuclei and moderate amounts of cytoplasm were interspersed among bands or sheets of parallel, elongated cells with thin, spindle-shaped, hyperchromatic nuclei. The myocardial schwannomas were typically less densely cellular and infiltrated amid, sometimes replacing, the cardiomyocytes. The cell types described for the endocardial neoplasms were both present, but in fewer numbers. In both subtypes of schwannomas, there was a minimal amount of cellular pleomorphism. In some larger neoplasms, Antoni type A and B patterns were present.

The Schwann cell hyperplasias were similar in appearance to the schwannomas, but were smaller and had less pleomorphism of the cells. In the case of the endocardial Schwann cell hyperplasia, there was no invasion of the myocardium.

## APPENDIX D HISTORICAL CONTROLS

**Table D1.** Incidence of astrocytoma, glioma, and/or oligodendroglioma in brains of male Harlan Sprague Dawley rats in NTP studies

| Chemical | First dose | N | Control incidence |
| --- | --- | --- | --- |
| Dibutylphthalate | 8/30/2010 | 49 | 4% |
| 2-Hydroxy-4-methoxybenzophenone | 11/8/2010 | 50 | 0% |
| p-Chloro-a,a,a-trifluorotoluene | 1/17/2011 | 50 | 4% |
| Di-(2-ethylhexyl)phthalate | 2/17/2011 | 50 | 8% |
| Di-(2-ethylhexyl)phthalate (perinatal) | 6/27/2011 | 50 | 0% |
| Tris (chloroisopropyl) phosphate | 12/12/2011 | 50 | 0% |
| Sodium tungstate | 12/23/2011 | 50 | 4% |
| Resveratrol | 5/7/2012 | 50 | 0% |
| Black cohosh | 7/2/2012 | 50 | 2% |
| Radiofrequency radiation (GSM/CDMA) | 9/16/2012 | 90 | 0% |

Historical control rate: 11/550 (2.0%)

**Table D2.** Incidence of schwannoma in the heart of male Harlan Sprague Dawley rats in NTP studies

| Chemical | First dose | N | Control incidence |
| --- | --- | --- | --- |
| Indole-3-carbinol | 3/14/2007 | 50 | 2% |
| Perfluorooctanoic acid | 6/19/2009 | 50 | 0% |
| Dietary zinc | 9/3/2009 | 50 | 0% |
| Dibutylphthalate | 8/30/2010 | 49 | 4% |
| 2-Hydroxy-4-methoxybenzophenone | 11/8/2010 | 50 | 2% |
| p-Chloro-a,a,a-trifluorotoluene | 1/17/2011 | 50 | 0% |
| Di-(2-ethylhexyl)phthalate | 2/17/2011 | 50 | 6% |
| Di-(2-ethylhexyl)phthalate (perinatal) | 6/27/2011 | 50 | 4% |
| Tris (chloroisopropyl) phosphate | 12/12/2011 | 50 | 0% |
| Sodium tungstate | 12/23/2011 | 50 | 0% |
| Resveratrol | 5/7/2012 | 50 | 0% |
| Black Cohosh | 7/2/2012 | 50 | 0% |
| Radiofrequency radiation (GSM/CDMA) | 9/16/2012 | 90 | 0% |

Historical control rate: 9/699 (1.30%)

## APPENDIX E TIME ON STUDY TO APPEARANCE OF TUMORS

**Malignant Glioma**

Malignant Glioma
| SAR (W/kg) | Animal ID number | Time on study (weeks) |
| --- | --- | --- |
| GSM-modulated exposed males |  |  |
| 1.5 | 717 | 105 |
|  | 735 | 102 |
|  | 786 | 104 |
| 3.0 | 924 | 101 |
|  | 943 | 105 |
|  | 1014 | 93 |
| 6.0 | 1135 | 104 |
|  | 1137 | 102 |
| CDMA-modulated exposed males |  |  |
| 6.0 | 1795 | 105 |
|  | 1799 | 104 |
|  | 1852 | 105 |
| GSM-modulated exposed females |  |  |
| 6.0 | 1246 | 96 |
| CDMA-modulated exposed females |  |  |
| 1.5 | 1463 | 105 |
|  | 1474 | 105 |
|  | 1523 | 550 |

**Time to Malignant Schwannoma in Heart**

Time to Malignant Schwannoma in Heart
| SAR (W/kg) | Animal ID number | Length of survival (weeks) |
| --- | --- | --- |
| GSM-modulated exposed males |  |  |
| 1.5 | 758 | 104 |
|  | 801 | 105 |
| 3.0 | 931 | 105 |
| 6.0 | 1149 | 83 |
|  | 1155 | 105 |
|  | 1187 | 104 |
|  | 1206 | 104 |
|  | 1230 | 91 |
| CDMA-modulated exposed males |  |  |
| 1.5 | 1364 | 105 |
|  | 1352 | 105 |
| 3.0 | 1559 | 92 |
|  | 1617 | 105 |
|  | 1622 | 104 |
| 6.0 | 1801 | 76 |
|  | 1821 | 70 |
|  | 1829 | 104 |
|  | 1833 | 89 |
|  | 1849 | 104 |
|  | 1860 | 105 |
| GSM-modulated exposed females |  |  |
| 3.0 | 1037 | 105 |
|  | 1077 | 83 |
| CDMA-modulated exposed females |  |  |
| 1.5 | 1461 | 106 |
|  | 1480 | 93 |
| 6.0 | 1888 | 105 |
|  | 1965 | 106 |

## APPENDIX F REVIEWERS’ COMMENTS

National Toxicology Program

Peer Review Charge and Summary Comments

<u>Purpose</u>: To provide independent peer review of an initial draft of this partial report. The peer reviewers were blind to the test agents under study. Introductory materials on RFR and details of the methods dealing with the field generation and animal housing were redacted from the version sent to the reviewers. The reviewers were provided a study data package, also blinded to test agents, containing basic in life study information such as body weight and survival curves and information concerning the generation of pups from the *in utero* exposures.

<u>Report Title</u>: Draft Report of Partial Findings from the National Toxicology Program Carcinogenesis Studies of Test Articles A and B (and associated Study Data Package)

<u>Reviewers’ Names</u>:

> David Dorman, D.V.M., Ph.D., North Carolina State University
>
> Russell Cattley, D.V.M., Ph.D., Auburn University
>
> Michael Pino, D.V.M., Ph.D., Pathology consultant

<u>Charge</u>: To peer review the draft report and comment on whether the scientific evidence supports NTP’s conclusion(s) for the study findings.

1. Scientific criticisms:
  a. Please comment on whether the information presented in the draft report, including presentation of data in any tables, is clearly and objectively presented. Please suggest any improvements. All three reviewers found the results to be clearly and objectively presented, although there were suggestions to provide historical control information for brain and heart lesions for female Harlan Sprague Dawley rats, clarify statements about the specific statistical tests used and the presence or lack of statistical significance of the brain gliomas in the Results, and expand the conclusions statements to clarify the basis for the conclusions.
  b. Please comment on whether NTP’s scientific interpretations of the data are objective and reasonable. Please explain why or why not. The reviewers stated that the NTP had performed an adequate and objective peer review of the pathology data, and the statistical approaches used were consistent with other NTP studies. The methods were described as objective and reasonable. The interpretations of the data, including the limitations, were also reasonable and objective. One reviewer found the data on schwannomas of the heart to be more compelling with respect to an association with treatment than the brain gliomas. This reviewer summarized the findings as:

> “In the heart the evidence for a carcinogenic effect can be based on 1) the presence of the tumors in all six of the test article groups versus none in the controls 2) the statistically significant trend for schwannomas with both compounds and the statistically significant increase in incidence in the 4X (top) dose for test article B; 3) the fact that the incidence of the tumors in both 4X dose groups approaches or exceeds the high end of the historical control range; and 4) the tumors in the 4X group of test article B are accompanied by a higher incidence of Schwann cell hyperplasia. Using the NTP’s guide for levels of evidence for carcinogenic activity, I would consider the heart schwannomas as ‘Some Evidence’ of carcinogenic activity.
>
> The proliferative lesions in the brain are more difficult to interpret because 1) their low incidence that was well within the historical control range, 2) lack of clear dose response; and 3) lack of statistical significance (except for the significant exposure-dependent trend for test article B. … However, the presence of malignant gliomas and/or foci of glial cell hyperplasia in 5 of 6 test article groups for both sexes vs none in controls of either sex is suggestive of a test article effect. …I would consider the malignant gliomas as ‘Equivocal Evidence’ of carcinogenic activity.”

2. Please identify any Information that should be added or deleted: One reviewer suggested that more information be given on the time when tumors were observed (e.g., at terminal necropsy, or early in the study) to help assess the possible impact of the decreased survival times in the control animals on tumor incidence. This reviewer also suggested a discussion of how the survival of control male rats in this study compared to the historical control data. There was also concern that the diagnostic criteria developed by the PWG and used in the current study would impact the historical control incidence rates reported in Table D.
3. The scientific evidence supports NTP’s conclusion(s) for the study findings: The NTP’s overall draft conclusion was as follows: “Under the conditions of these studies, the observed hyperplastic lesions and neoplasms outlined in this partial report are considered likely the result of exposures to test article A and test article B. The findings in the heart were statistically stronger than the findings in the brain.” The reviewers had the option of agreeing, agreeing in principle, or disagreeing with the draft conclusions. All three reviewers agreed in principle, reiterating issues discussed above.

## APPENDIX G NIH 1 REVIEWERS’ COMMENTS

<u>Purpose</u>: To provide independent peer review of the pathology diagnoses and statistical evaluation of the partial findings from NTP’s studies. Background materials included the draft NTP report, introductory materials on RFR, and details on the methods dealing with the field generation and statistical analyses references and guidance. The reviewers were provided a stud data package, containing basic in life study information such as body weight and survival curve information concerning the generation of pups from the *in utero* exposures, and raw pathology data.

<u>Reviewers’ Names</u>:

> Diana C. Haines, D.V.M., Frederick National Laboratory
>
> Michael S. Lauer, M.D., Office of Extramural Research, NIH
>
> Maxwell P. Lee, Ph.D., Laboratory of Cancer Biology and Genetics, NCI,
>
> Aleksandra M. Michalowski, M.Sc., Ph.D., Laboratory of Cancer Biology and Genetics, NCI
>
> R. Mark Simpson, D.V.M., Ph.D., Laboratory of Cancer Biology and Genetics, NCI
>
> [Sixth reviewer’s name and comments are withheld.]

<u>Charge</u>: To peer review the draft report, statistical analyses, and pathology data and comment o whether the scientific evidence supports NTP’s conclusion(s) for the study findings.

Reviewer’s comments and NTP responses to the comments are provided.

- Appendix G1: Reviewers’ comments
- Appendix G2: NTP’s responses to NIH reviewers’ comments

## Appendix G1 Reviewers’ Comments

Reviewer: Diana C. Haines, D.V.M., Frederick National Laboratory

April 5, 2016

Dr. Tabak,

I’ve always relied on experts, not myself, for statistical analysis, and so do not feel qualified to address the statistical methods used. My training and experience has been in veterinary pathology, including QA review of NTP studies, and serving on PWGs, so will give my opinion on the pathology interpretation (biological significance rather than statistical significance).

Having perused the 3 RFR Draft Report and the raw data, all appears to be in order, including QA of the histopathology (technique) as well as PWG review (diagnosis). Looking at the data, I agree with the report’s conclusion: *Under the conditions of these studies, the hyperplastic lesions and neoplasms observed in male rats are considered likely the result of exposures to GSM-and CDMA-modulated RFR. The findings in the heart were statistically stronger than the findings in the brain.* But note, it is “considered likely” not “definitely is”.

There may be also several caveats relating to “under the conditions of these studies”, including how well the conditions recapitulate actual human exposure: whole body exposure from in utero to old age; 18.5 hours/day (10 min on/10 min off, for total of 9hr actual exposure); and dose^A^. I’m not a physicist, so have to presume experts analyzed and accepted concept of the reverberation chamber, including “doses”^A^, as being relevant to human exposure.

^A^ Dosimetric Assessment paper: “As could be expected in a study following NTP protocols, the exposure levels for the rodents in this project exceed the limits for the wbSAR and psSAR defined in the IEEE Std C95.1-2005 safety standard for human exposure to mobile phone radiation. In the low dose exposure group the exposure level in the organs exceeds or is close to the localized SAR limit for the general public, except for a few low-water content tissues. More specifically, the psSAR over 1 g in the human head, is limited by the safety standards to <2W/kg, whereas, in the low dose rodents the SAR averaged over the whole brain is >2.4 W/kg for mice, and >1.3 W/kg for rats, hence similar to the limit. Furthermore, the psSAR and oSAR have larger uncertainty compared to the wbSAR. Deviations of the exposure level from the target dose, especially during the early exposure period, should be carefully evaluated in the interpretation of the final biological studies.

Results from the companion mouse study will hopefully add some insight.

**Diana Copeland Haines, DVM**

Diplomate, American College of Veterinary Pathologists

Senior Staff Pathologist, Pathology Section

Pathology/Histotechnology Laboratory

Leidos Biomedical Research, Inc.

Frederick National Laboratory for Cancer Research

P.O. Box B, Frederick, MD 21702

Phone: 301-846-5921 Fax: 301-846-1953

http://ncifrederick.cancer.gov/rtp/lasp/phl/

Reviewer: Michael S. Lauer, M.D., Office of Extramural Research, NIH

Michael S Lauer, MD (OER)

Review of NTP paper: “Report of Partial Findings from the National Toxicology Program Carcinogenesis Studies of Cell Phone Radiofrequency Radiation (Whole Body Exposures)”

March 20, 2016

Summary of findings:

This is a partial report, a report which is presumably part of a larger set of studies involving 2 species (mice and rats), 2 sexes (male, female), and multiple tissue types, all based on 90-week studies of two different types (GSM and CDMA) of cell phone radiofrequency radiation (RFR). In this partial report, we are given findings regarding brain gliomas and heart schwannomas in male and female Harlan Sprague Dawley rats which were exposed exposed to control or 3 different levels (1.5, 3.0, 6.0) of two types (GSM and CDMA) of RFR. There were 90 rats in each group. Using the poly-3 test with the Bieler-Williams variance adjustment, the authors found a statistically significant increase in the rate of brain gliomas in males exposed to CDMA RFR. Using the poly-6 test, the authors found a statistically significant increase in the rates of heart schwannomas in males exposed to GSM and CDMA. There were no statistically significant differences in rates of gliomas or schwannomas in females; also there was no statistically significant increase in rates of gliomas in males exposed to GSM RFR.

Comments:

1. Why aren’t we being told, at least at a high level, of the results of other experiments (i.e., male and female mice, tissues other than heart and brain, tumors other than glioma and schwannoma)? Given the multiple comparisons inherent in this kind of work (see pages 27-30 and Table 13 of the FDA guidance document), there is a high risk of false positive discoveries. In the absence of knowing other findings, we must worry about selective reporting bias.
2. I was able to reproduce the authors’ positive P-value findings (see Appendix 1, R code) using the <u>MCPAN R package</u>. However, I’m getting slightly different values for adjusted denominators (also in Appendix 1).
3. I was able to reproduce the authors’ findings of longer survival with RFR (see Appendix 1, R code).
4. I have a number of questions about the study design:

a. Were control rats selected in utero like the exposed rats were?
b. Were pregnant dams assigned to different groups by formal randomization? If not, why not?
c. Why were pups in the same litter included? Did the authors take any steps in their analyses to account for the resulting absence of i.i.d?
d. The authors state that at most 3 pups were chosen per litter. How were the 3pups chosen (and the others presumably not used for this experiment)? Were the 3 pups that were chosen selected by formal randomization? If not, why not?
e. Were all analyses based on the intent-to-treat principle? Were there any crossovers? Were all rats accounted for by the end of the experiment and were all rats who started in the experiment included in the final analyses?
f. Blinding: The authors state that “All PWG reviewer were conducted blinded with respect to treatment group,” but in the very next phrase write “only identifying the test articles as ‘test agent A’ or ‘test agent B.’” Why was this information(test agent A or B) given? The blinding was not complete.
5. Sample size:

a. Did the authors perform a prospective (that is before initiation of the work)sample size calculation? If so, what were the prior assumptions? In other words,why did the authors choose to study 90 rats in each group and why did they set the maximum duration to 90 weeks (instead of 104 weeks)?
b. I used a <u>publicly available</u> simulation package^1^ to calculate the study power for male rats based on the following (see Appendix 2, power calculation simulationstudies):

i. Control tumor rate of ~1.5%.
ii. Risk ratio 2.5 in the group receiving the highest dose
iii. 2-sided Alpha = 0.005 (based on Table 13 of the FDA guidance document). Note this low alpha of 0.005 for poly-k trend tests is recommended to minimize the risk of false positive discoveries.
iv. Sample size of 90 for each group with one planned sacrifice.
v. Low lethality with lethality parameters set according to study duration and Weibull shape parameter (see Table 3 of Moon et al^1^). When I re-ran the simulations using intermediate lethality, results were not materially changed.
vi. Study duration 90 weeks
vii. 5000 simulations
viii. Note - I used dose levels of 0,1,2, and 4 because I was unable to adjust these on the web site (despite trying 3 different browsers).
c. Based on these inputs, the recommendations in Table 13 of the FDA guidance document, and a sample size of 90 rats in each group, I find very low power(<5%, see Appendix 2). Even allowing for a risk ratio of 5.0 (a level that is clinically unlikely), the power for 2-sided alpha=0.005, k=3 and low lethality is only ~14% (see Appendix 2).
d. The low power implies that there is a high risk of false positive findings^2^,especially since the epidemiological literature questions the purported association between cell phone exposure and cancer.^3^
6. Summary: I am unable to accept the authors’conclusions:

a. We need to know all other findings of these experiments (mice, other tumor types) given the risk of false positive findings and reporting bias. It would be helpful to have a copy of the authors’ statistical code.
b. We need to know whether randomization was employed to assign dams to specific groups (control and intervention).
c. We need to know whether randomization was employed to determine which pups from each litter were chosen for continued participation in the experiment.
d. We need to know whether there was a formal power/sample size calculation performed prior to initiation of the experiment. If not, why not? If yes, we need to see the details. In particular, we need to know whether the authors followed the recommendations of the FDA guidance document (in particular Table 13).
e. I suspect that this experiment is substantially underpowered and that the few positive results found reflect false positive findings.^2^ The higher survival with RFR, along with the prior epidemiological literature, leaves me even more skeptical of the authors’ claims.

## Appendix 1: Attempted replication of positive findings

\# Review of NTP paper on cell phone RFR and certain cancers

\# Attempt to reproduce the positive findings

\# Data from Larry Tabak

\# Code by Mike Lauer

setwd("~/Desktop/Files to save")

library(MCPAN)

library(rms)

library(Hmisc)

\# Read in CDMA NTP data

CDMA <- read.csv("~/Desktop/Files to save/NTP CDMA Raw Tumor Data.csv")

\# Survival and treatment group, adjusting for sex, by Cox proportional hazards

CDMA$status<-1

CDMA$S<-Surv(CDMA$Removal.Day, CDMA$status)

f<-cph(S~Treatment+Sex, data=CDMA)

f

\# Survival greater (better) for 3.0W, P=0.0157, for 6.0W, P=0.0260

\# Table 1 – Poly-3 test for malignant glioma in males CDMA

males_CDMA<-subset(CDMA, Sex==’M’)

poly3test(time=males_CDMA$Removal.Day, status=males_CDMA$Brain.Glioma.Malignant, f=males_CDMA$Dose, k=3, type=’Williams’, method=’BW’, alternative=’greater’)

\# P=0.039

poly3ci(time=males_CDMA$Removal.Day, status=males_CDMA$Brain.Glioma.Malignant, f=males_CDMA$Dose, k=3, type=’Williams’, method=’BW, alternative=’greater’)

Call result:

Sample estimates, using poly-3-adjustment

|  | 0 | 1.5 | 3 | 6 |
| --- | --- | --- | --- | --- |
| x | 0.0000 | 0.0000 | 0.0000 | 3.0000 |
| n | 90.0000 | 90.0000 | 90.0000 | 90.0000 |
| adjusted n | 63.8258 | 72.3688 | 76.6821 | 64.8154 |
| adjusted estimate | 0.0000 | 0.0000 | 0.0000 | 0.0463 |

\# Table 3 - Poly-6 test for malignant Schwannoma in males CDMA

poly3test(time=males_CDMA$Removal.Day, status=males_CDMA$Heart.Schwannoma.Malignant, f=males_CDMA$Dose, k=6, type=’Williams’, method=’BW’, alternative=’greater’)

\# P=0.0005

poly3ci(time=males_CDMA$Removal.Day, status=males_CDMA$Heart.Schwannoma.Malignant,f=males_CDMA$Dose, k=3,type=’Williams’, method=’BW’)

Call result:

Sample estimates, using poly-3-adjustment

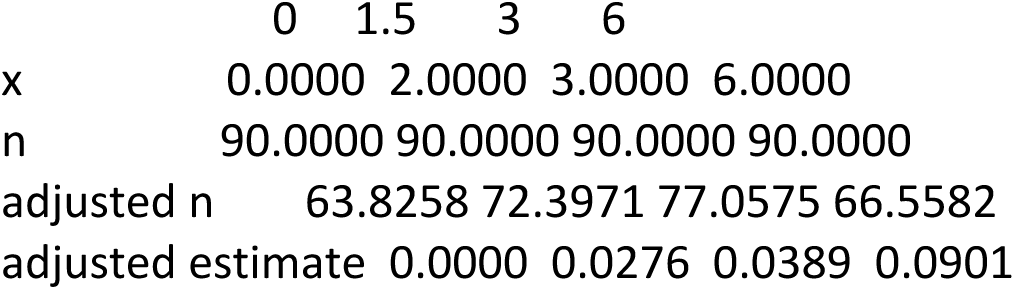

|  | 0 | 1.5 | 3 | 6 |
| --- | --- | --- | --- | --- |
| x | 0.0000 | 2.0000 | 3.0000 | 6.0000 |
| n | 90.0000 | 90.0000 | 90.0000 | 90.0000 |
| adjusted n | 63.8258 | 72.3971 | 77.0575 | 66.5582 |
| adjusted estimate | 0.0000 | 0.0276 | 0.0389 | 0.0901 |

\# Read in GSM NTP data

GSM <- read.csv("~/Desktop/Files to save/NTP GSM Raw Tumor data.csv")

\# Survival and treatment group, adjusting for sex, by Cox proportional hazards

GSM$status<-1

GSM$S<-Surv(GSM$Removal.Day, GSM$status)

f<-cph(S~Treatment+Sex, data=GSM)

f

\# Survival greater (better) for 6.0W, P=0.0048

males_GSM<-subset(GSM, Sex==’M’)

\# Table 3 – Poly-6 test for malignant Schwannomas in males GSM

poly3test(time=males_GSM$Removal.Day, status=males_GSM$Heart.Schwannoma.Malignant, f=males_CDMA$Dose, k=6, type=’Williams’, method=’BW’, alternative=’greater’)

\# P=0.004

poly3ci(time=males_GSM$Removal.Day, status=males_GSM$Heart.Schwannoma.Malignant, f=males_CDMA$Dose, k=3, type=’Williams’, method=’BW’, alternative=’greater’)

Call result:

Sample estimates, using poly-3-adjustment

|  | 0 | 1.5 | 3 | 6 |
| --- | --- | --- | --- | --- |
| x | 0.0000 | 2.0000 | 1.0000 | 5.0000 |
| n | 90.0000 | 90.0000 | 90.0000 | 90.0000 |
| adjusted n | 63.8258 | 73.1547 | 76.1127 | 77.0723 |
| adjusted estimate | 0.0000 | 0.0273 | 0.0131 | 0.0649 |

## Appendix 2: Simulations for power calculations

Power Simulations for NTP Cell Phone RFR paper (from https://biostatistics.mdanderson.org/acss/Login.aspx and https://www.jstatsoft.org/article/view/v007i13)^1^

Michael Lauer, MD (OER)

March 19, 2016

> 1) For malignant gliomas (Table 1), P = 0.005, HR = 2.5, k=3

The University of Texas M. D. Anderson Cancer Center Sample Size and Power Estimation for Animal Carcinogenicity Studies

*** Input Parameters ***

Selected Seed = 3000

Number of Groups = 4

Dose metric of each group:

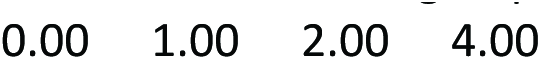

Number of animals in each group

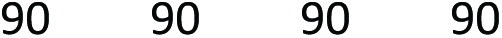

Number of sacrifices including a terminal sacrifice = 1

Sacrifice time points in weeks:

Study duration = 90 weeks

Number of INTERIM sacrificed animals in each interval:

Background tumor onset probability at the end of the study = 0.01

Tumor onset distribution assumed: Weibull with a shape parameter 3.00

Hazard ratio(s) of dose vs. control group

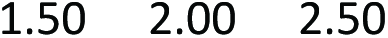

Competing Risks Survival Rate (CRSR) for each group:

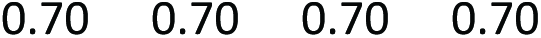

Tumor lethality parameter entered = 23.00

Level of the test = 0.01

One-sided or two-sided test = 2 sided test

Number of simulation runs = 5000

*** Simulation Results ***

dose group 0:

average tumor rate = 0.0149

average competing risks survival rate = 0.6990

average lethality = 0.0816

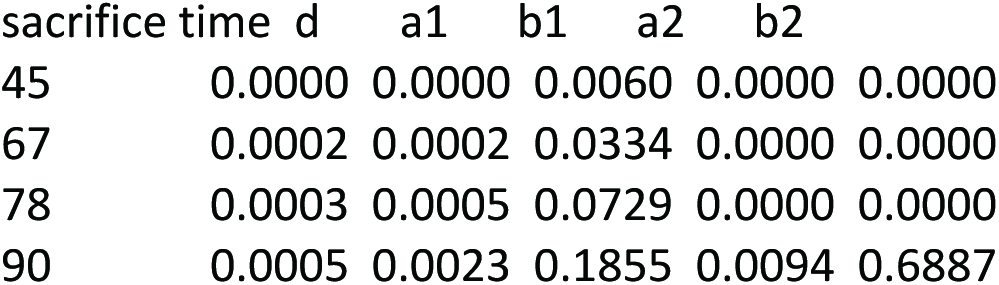

dose group 1:

average tumor rate = 0.0225

average competing risks survival rate = 0.7000

average lethality = 0.0784

| sacrifice time | d | a1 | b1 | a2 | b2 |
| --- | --- | --- | --- | --- | --- |
| 45 | 0.0001 | 0.0000 | 0.0059 | 0.0000 | 0.0000 |
| 67 | 0.0003 | 0.0002 | 0.0325 | 0.0000 | 0.0000 |
| 78 | 0.0004 | 0.0008 | 0.0720 | 0.0000 | 0.0000 |
| 90 | 0.0007 | 0.0034 | 0.1851 | 0.0145 | 0.6842 |

dose group 2:

average tumor rate = 0.0297

average competing risks survival rate = 0.6997

average lethality = 0.0772

| sacrifice time | d | a1 | b1 | a2 | b2 |
| --- | --- | --- | --- | --- | --- |
| 45 | 0.0001 | 0.0000 | 0.0059 | 0.0000 | 0.0000 |
| 67 | 0.0004 | 0.0003 | 0.0331 | 0.0000 | 0.0000 |
| 78 | 0.0005 | 0.0012 | 0.0721 | 0.0000 | 0.0000 |
| 90 | 0.0010 | 0.0045 | 0.1829 | 0.0191 | 0.6790 |

dose group 3:

average tumor rate = 0.0366

average competing risks survival rate = 0.7007

average lethality = 0.0772

| sacrifice time | d | a1 | b1 | a2 | b2 |
| --- | --- | --- | --- | --- | --- |
| 45 | 0.0001 | 0.0000 | 0.0059 | 0.0000 | 0.0000 |
| 67 | 0.0005 | 0.0003 | 0.0330 | 0.0000 | 0.0000 |
| 78 | 0.0006 | 0.0013 | 0.0716 | 0.0000 | 0.0000 |
| 90 | 0.0012 | 0.0054 | 0.1812 | 0.0238 | 0.6749 |

Positive Trend (Power): 0.0238

> 2) For malignant Schwannomas (Table 3), P = 0.005, HR = 2.5, k=6

The University of Texas M. D. Anderson Cancer Center

Sample Size and Power Estimation for Animal Carcinogenicity Studies

*** Input Parameters ***

Selected Seed = 3000

Number of Groups = 4

Dose metric of each group:

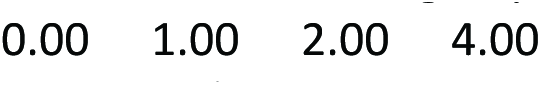

Number of animals in each group

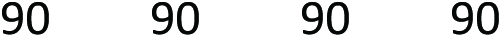

Number of sacrifices including a terminal sacrifice = 1

Sacrifice time points in weeks:

Study duration = 90 weeks

Number of INTERIM sacrificed animals in each interval:

Background tumor onset probability at the end of the study = 0.01

Tumor onset distribution assumed: Weibull with a shape parameter 6.00

Hazard ratio(s) of dose vs. control group

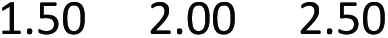

Competing Risks Survival Rate (CRSR) for each group:

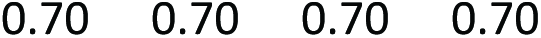

Tumor lethality parameter entered = 45.00

Level of the test = 0.01

One-sided or two-sided test = 2 sided test

Number of simulation runs = 5000

*** Simulation Results ***

dose group 0:

average tumor rate = 0.0149

average competing risks survival rate = 0.6990

average lethality = 0.0631

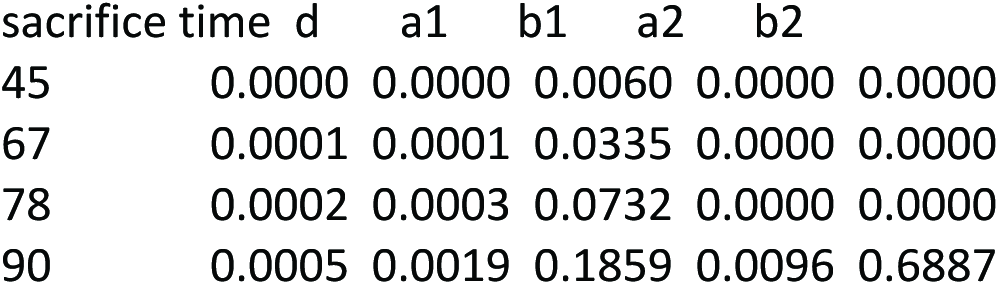

| sacrifice time | d | a1 | b1 | a2 | b2 |
| --- | --- | --- | --- | --- | --- |
| 45 | 0.0000 | 0.0000 | 0.0060 | 0.0000 | 0.0000 |
| 67 | 0.0001 | 0.0001 | 0.0335 | 0.0000 | 0.0000 |
| 78 | 0.0002 | 0.0003 | 0.0732 | 0.0000 | 0.0000 |
| 90 | 0.0005 | 0.0019 | 0.1859 | 0.0096 | 0.6887 |

dose group 1:

average tumor rate = 0.0225

average competing risks survival rate = 0.7000

average lethality = 0.0602

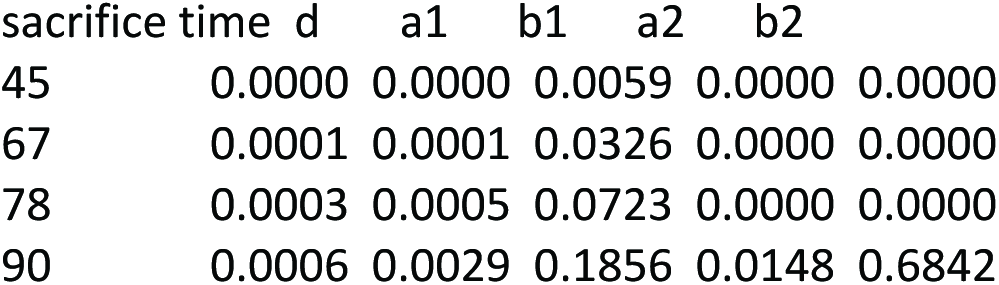

dose group 2:

average tumor rate = 0.0297

average competing risks survival rate = 0.6997

average lethality = 0.0582

| sacrifice time | d | a1 | b1 | a2 | b2 |
| --- | --- | --- | --- | --- | --- |
| 45 | 0.0000 | 0.0000 | 0.0059 | 0.0000 | 0.0000 |
| 67 | 0.0002 | 0.0001 | 0.0333 | 0.0000 | 0.0000 |
| 78 | 0.0004 | 0.0007 | 0.0726 | 0.0000 | 0.0000 |
| 90 | 0.0009 | 0.0038 | 0.1837 | 0.0195 | 0.6790 |

dose group 3:

average tumor rate = 0.0366

average competing risks survival rate = 0.7007

average lethality = 0.0588

| sacrifice time | d | a1 | b1 | a2 | b2 |
| --- | --- | --- | --- | --- | --- |
| 45 | 0.0000 | 0.0000 | 0.0059 | 0.0000 | 0.0000 |
| 67 | 0.0003 | 0.0001 | 0.0332 | 0.0000 | 0.0000 |
| 78 | 0.0005 | 0.0007 | 0.0722 | 0.0000 | 0.0000 |
| 90 | 0.0011 | 0.0046 | 0.1821 | 0.0243 | 0.6749 |

Positive Trend (Power): 0.0230

> 3) For further consideration, P = 0.005, HR = 5, k=3

The University of Texas M. D. Anderson Cancer Center

Sample Size and Power Estimation for Animal Carcinogenicity Studies

*** Input Parameters ***

Selected Seed = 3000

Number of Groups = 4

Dose metric of each group:

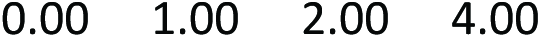

Number of animals in each group

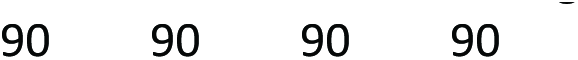

Number of sacrifices including a terminal sacrifice = 1

Sacrifice time points in weeks:

Study duration = 90 weeks

Number of INTERIM sacrificed animals in each interval:

Background tumor onset probability at the end of the study = 0.01

Tumor onset distribution assumed: Weibull with a shape parameter 3.00

Hazard ratio(s) of dose vs. control group

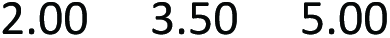

Competing Risks Survival Rate (CRSR) for each group:

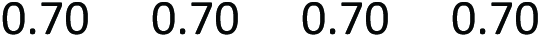

Tumor lethality parameter entered = 23.00

Level of the test = 0.01

One-sided or two-sided test = 2 sided test

Number of simulation runs = 5000

*** Simulation Results ***

dose group 0:

average tumor rate = 0.0149

average competing risks survival rate = 0.6990

average lethality = 0.0816

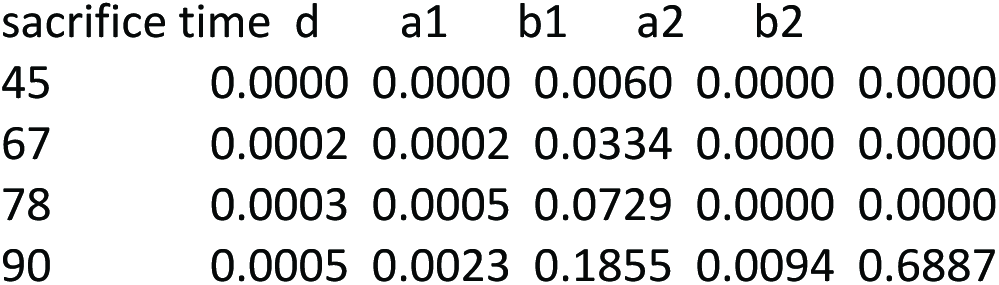

dose group 1:

average tumor rate = 0.0301

average competing risks survival rate = 0.7000

average lethality = 0.0743

| sacrifice time | d | a1 | b1 | a2 | b2 |
| --- | --- | --- | --- | --- | --- |
| 45 | 0.0001 | 0.0000 | 0.0059 | 0.0000 | 0.0000 |
| 67 | 0.0004 | 0.0003 | 0.0324 | 0.0000 | 0.0000 |
| 78 | 0.0005 | 0.0011 | 0.0717 | 0.0000 | 0.0000 |
| 90 | 0.0009 | 0.0045 | 0.1839 | 0.0194 | 0.6789 |

dose group 2:

average tumor rate = 0.0515

average competing risks survival rate = 0.6997

average lethality = 0.0774

| sacrifice time | d | a1 | b1 | a2 | b2 |
| --- | --- | --- | --- | --- | --- |
| 45 | 0.0002 | 0.0000 | 0.0058 | 0.0000 | 0.0000 |
| 67 | 0.0007 | 0.0006 | 0.0328 | 0.0000 | 0.0000 |
| 78 | 0.0009 | 0.0020 | 0.0713 | 0.0000 | 0.0000 |
| 90 | 0.0017 | 0.0076 | 0.1795 | 0.0331 | 0.6638 |

dose group 3:

average tumor rate = 0.0727

average competing risks survival rate = 0.7007

average lethality = 0.0804

| sacrifice time | d | a1 | b1 | a2 | b2 |
| --- | --- | --- | --- | --- | --- |
| 45 | 0.0003 | 0.0000 | 0.0059 | 0.0000 | 0.0000 |
| 67 | 0.0010 | 0.0006 | 0.0327 | 0.0000 | 0.0000 |
| 78 | 0.0013 | 0.0028 | 0.0701 | 0.0000 | 0.0000 |
| 90 | 0.0025 | 0.0107 | 0.1755 | 0.0470 | 0.6496 |

Positive Trend (Power): 0.1420

4) For further consideration, same as in baseline (1) but with intermediate lethality

*** Input Parameters ***

Selected Seed = 3000

Number of Groups = 4

Dose metric of each group:

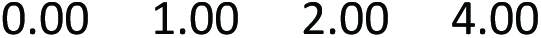

Number of animals in each group

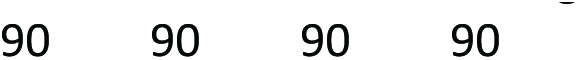

Number of sacrifices including a terminal sacrifice = 1

Sacrifice time points in weeks:

Study duration = 90 weeks

Number of INTERIM sacrificed animals in each interval:

Background tumor onset probability at the end of the study = 0.01

Tumor onset distribution assumed: Weibull with a shape parameter 3.00

Hazard ratio(s) of dose vs. control group

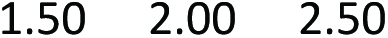

Competing Risks Survival Rate (CRSR) for each group:

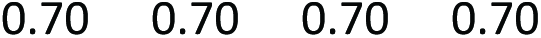

Tumor lethality parameter entered = 225.00

Level of the test = 0.01

One-sided or two-sided test = 2 sided test

Number of simulation runs = 5000

*** Simulation Results ***

dose group 0:

average tumor rate = 0.0149

average competing risks survival rate = 0.6990

average lethality = 0.3936

| sacrifice time | d | a1 | b1 | a2 | b2 |
| --- | --- | --- | --- | --- | --- |
| 45 | 0.0004 | 0.0000 | 0.0060 | 0.0000 | 0.0000 |
| 67 | 0.0014 | 0.0001 | 0.0334 | 0.0000 | 0.0000 |
| 78 | 0.0014 | 0.0004 | 0.0729 | 0.0000 | 0.0000 |
| 90 | 0.0019 | 0.0015 | 0.1855 | 0.0063 | 0.6887 |

dose group 1:

average tumor rate = 0.0225

average competing risks survival rate = 0.7000

average lethality = 0.3852

| sacrifice time | d | a1 | b1 | a2 | b2 |
| --- | --- | --- | --- | --- | --- |
| 45 | 0.0006 | 0.0000 | 0.0059 | 0.0000 | 0.0000 |
| 67 | 0.0022 | 0.0001 | 0.0325 | 0.0000 | 0.0000 |
| 78 | 0.0020 | 0.0006 | 0.0720 | 0.0000 | 0.0000 |
| 90 | 0.0029 | 0.0023 | 0.1851 | 0.0097 | 0.6842 |

dose group 2:

average tumor rate = 0.0297

average competing risks survival rate = 0.6997

average lethality = 0.3839

| sacrifice time | d | a1 | b1 | a2 | b2 |
| --- | --- | --- | --- | --- | --- |
| 45 | 0.0008 | 0.0000 | 0.0059 | 0.0000 | 0.0000 |
| 67 | 0.0029 | 0.0003 | 0.0331 | 0.0000 | 0.0000 |
| 78 | 0.0027 | 0.0008 | 0.0721 | 0.0000 | 0.0000 |
| 90 | 0.0039 | 0.0031 | 0.1829 | 0.0127 | 0.6790 |

dose group 3:

average tumor rate = 0.0366

average competing risks survival rate = 0.7007

average lethality = 0.3897

| sacrifice time | d | a1 | b1 | a2 | b2 |
| --- | --- | --- | --- | --- | --- |
| 45 | 0.0009 | 0.0000 | 0.0059 | 0.0000 | 0.0000 |
| 67 | 0.0037 | 0.0003 | 0.0330 | 0.0000 | 0.0000 |
| 78 | 0.0033 | 0.0009 | 0.0716 | 0.0000 | 0.0000 |
| 90 | 0.0048 | 0.0037 | 0.1812 | 0.0157 | 0.6749 |

Positive Trend (Power): 0.0219

Reviewer: Maxwell P. Lee, Ph.D., Laboratory of Cancer Biology and Genetics, NCI

I think the study was well designed and the analyses and results were clearly presented.

My main concern is the control data. Since the main finding was the increased incidence rates of heart schwannomas and brain gliomas in male Harlan Sprague Dawley rats exposed to GSM-or CDMA-modulated cell phone RFR, my analyses and evaluation below were focused on the male rats.

My concern regarding the control data came from the following two considerations. First, we need to consider sample variation. The incidence rates of the current controls for brain gliomas and heart schwannomas were 0. However, the historical controls were 1.67% for gliomas (range 0-8%) and 1.30% for schwannomas (0-6%). Given that there were substantial variations among the historical controls and the concurrent control is at the lowest end of the range, it is important to evaluate how different estimates of control incidence rates may impact the results of analyses. Supplementary Table S1 shows that for gliomas with 1.7% incidence rate we have 40%, 37%, 17%, and 6% of chance to observe 0 tumor, 1 tumor, 2 tumors, and greater than 2 tumors, respectively; heart schwannomas has similar distribution. Given the low incidence rate and moderate sample size of the control, even after observing 0 tumor in the current study, the true incidence rate may be higher than 0. If we were repeating the experiment, we may see some control studies have 1 or more tumors. Second, it is puzzling why the control had short survival rate. Given that most of the gliomas and heart schwannomas are late-developing tumors, it is possible that if the controls were living longer some tumors might develop. Although the use of poly-3 (or poly-6) test intended to adjust the number of rats used in the study, it is still important to re-evaluate the analysis by considering the incidence rate in controls not being 0.

Therefore, I have performed the analyses using the original data as well as the data modified by adding 1 tumor to the control. I implemented the poly-3 (or poly-6) trend test in R using the formula described in the file, Poly3 correction factor[1].docx.

The results are summarized in Table 1 for brain gliomas.

**Table 1.**
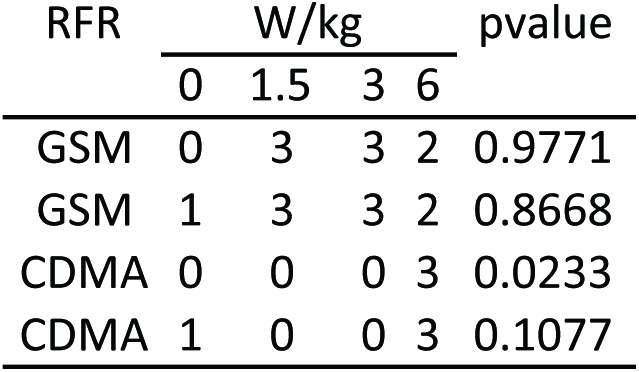
**Incidence of brain gliomas in male rats exposed to GSM-or CDMA-modulated RFR, comparing control data with 0 vs. 1 tumor.**

| RFR | W/kg |  |  |  | pvalue |
| --- | --- | --- | --- | --- | --- |
|  | 0 | 1.5 | 3 | 6 |  |
| GSM | 0 | 3 | 3 | 2 | 0.9771 |
| GSM | 1 | 3 | 3 | 2 | 0.8668 |
| CDMA | 0 | 0 | 0 | 3 | 0.0233 |
| CDMA | 1 | 0 | 0 | 3 | 0.1077 |

Poly-6 adjusted rates were used in the chi-square trend test. The 1^st^ and 3^rd^ rows correspond to the original data with 0 tumor observed in the control group (The numbers in Table 1 here are identical to those in Table 1 in the original report). The test is significant for CDMA exposures (pvalue = 0.0233). However, it is not significant after adding 1 tumor to the control group (pvalue = 0.1077, the 4^th^ row).

Similar analysis was performed for heart schwannomas. The results are summarized in Table 2.

**Table 2.** **Incidence of heart schwannomas in male rats exposed to GSM-or CDMA-modulated RFR, comparing control data with 0 vs. 1 tumor.**

| RFR | W/kg |  |  |  | pvalue |
| --- | --- | --- | --- | --- | --- |
|  | 0 | 1.5 | 3 | 6 |  |
| GSM | 0 | 2 | 1 | 5 | 0.0431 |
| GSM | 1 | 2 | 1 | 5 | 0.1079 |
| CDMA | 0 | 2 | 3 | 6 | 0.0144 |
| CDMA | 1 | 2 | 3 | 6 | 0.0365 |

Poly-3 adjusted rates were used in the chi-square trend test. The 1^st^ and 3^rd^ rows correspond to the original data with 0 tumor observed in the control group (The numbers in Table 2 here are identical to those in Table 3 in the original report). The tests are significant for both GSM (pvalue = 0.0431) and CDMA (pvalue = 0.0144) exposures. However, only CDMA exposure remains significant after adding 1 tumor to the control group (pvalue = 0.0365, the 4^th^ row).

Since the incidence of heart schwannomas in the 6 W/kg males was significantly higher in CDMA exposed males than the control group in the original report, I also analyzed the impact of adding 1 tumor to the control group.

**Table 3.** **Incidence of heart schwannomas in male rats exposed to 6 W/kg CDMA-modulated RFR, comparing control data with 0 vs. 1 tumor.**

| RFR | W/kg |  | pvalue |
| --- | --- | --- | --- |
|  | 0 | 6 |  |
| CDMA | 0 | 6 | 0.0381 |
| CDMA | 1 | 6 | 0.0986 |

Poly-3 adjusted rates were used in the chi-square trend test. The 1^st^ row corresponds to the original data with 0 tumor observed in the control group. The test was significant for CDMA exposures (pvalue = 0.0381). However, it was not significant after adding 1 tumor to the control group (pvalue = 0.0986, the 2^nd^ row).

**Conclusions**

Increased incidence of heart schwannomas in male rats exposed to GSM-or CDMA-modulated RFR is statistically significant by the chi-square trend test. The evidence is better for CDMA exposure than GSM exposure. I think additional experiments are needed to assess if the incidence of brain gliomas in male rats exposed to GSM-or CDMA-modulated RFR is significantly higher than the control group or not.

My additional comments are summarized below.

1. I compared poly-3 adjusted number from Table 3 in the original report versus the poly-3 adjusted number that I calculated using the raw data from the excel files. Supplementary Figure S1 shows that these two sets of numbers agree with each other in general. This is in contrast to the comparison for poly-6 adjusted number from Table 1 in the original report versus the poly-6 adjusted number that I calculated using the raw data from the excel files (Supplementary Figure S2). In fact, the adjusted rat numbers from Table 1 and Table 3 of the original report look quite similar (Supplementary Figure S3). This suggests that the poly-3 adjusted number was used in the footnotes in both Table 1 and Table 3 in the original report.

2. I noted that in Table S2 the adjusted numbers in from.original.report and poly3 are identical at Dose 0 and 1.5 for both CDMA and GSM as well as at Dose 3 for GSM but differ slightly in the other treatment doses for heart schwannomas. One possible cause of the difference is that the version of the raw data in the excel files differs from that used to generate the original report. The second possibility is typo in the footnote in Table 3. I also generated Table S3 that has the poly-6 adjusted numbers for brain gliomas. The two sets of the poly-6 adjusted numbers are very different.

3. There are a couple of errors in the footnote of Table 3 in the original report. 2/74.05 (5%) should be 2/74.05 (2.7%). 3/78.67 (4%) should be 3/78.67 (3.8%).

## Supplementary Information

**Table S1.** **Expected percentage of observing different numbers of tumors in the controls based on binomial distribution.**

|  | 0 tumor | 1 tumor | 2 tumors | >2 tumors |
| --- | --- | --- | --- | --- |
| control for glioma | 40% | 37% | 17% | 6% |
| control for heart schwannoma | 43% | 37% | 15% | 5% |

The percentage was calculated with 1.7% historical control rate for male rats (gliomas) and with poly-6 adjusted animal number, 53. Similarly, the percentage was calculated with 1.3% historical control rate for male (heart schwannoma) and with poly-3 adjusted animal number, 65.

**Table S2.** **The poly-3 adjusted rat numbers in Table 3 in the original report and those calculated from the raw data.**

| RFR | Dose | from.original.report | poly3 |
| --- | --- | --- | --- |
| CDMA | 0 | 65.47 | 65.47 |
| CDMA | 1.5 | 74.05 | 74.05 |
| CDMA | 3 | 78.67 | 78.35 |
| CDMA | 6 | 67.94 | 66.24 |
| GSM | 0 | 65.47 | 65.47 |
| GSM | 1.5 | 74.87 | 74.87 |
| GSM | 3 | 77.89 | 77.89 |
| GSM | 6 | 78.48 | 77.66 |

The numbers in from.original.report refers to the poly-3 adjusted rat number from Table 3 in the original report. The numbers in poly3 refers to the poly-3 adjusted rat numbers that I calculated from the raw data for heart schwannoma.

**Table S3.** **The poly-6 adjusted rat numbers in Table 1 in the original report and those calculated from the raw data.**

| RFR | Dose | from.original.report | poly6 |
| --- | --- | --- | --- |
| CDMA | 0 | 65.47 | 53.48 |
| CDMA | 1.5 | 74.05 | 65.94 |
| CDMA | 3 | 78.35 | 73.08 |
| CDMA | 6 | 66.24 | 57.5 |
| GSM | 0 | 65.47 | 53.48 |
| GSM | 1.5 | 74.93 | 67.84 |
| GSM | 3 | 78.27 | 71.43 |
| GSM | 6 | 77.1 | 72.55 |

The numbers in from.original.report refers to the poly-6 adjusted rat number from Table 1 in the original report. The numbers in poly6 refers to the poly-6 adjusted rat numbers that I calculated from the raw data for brain gliomas.

**Figure S1.**
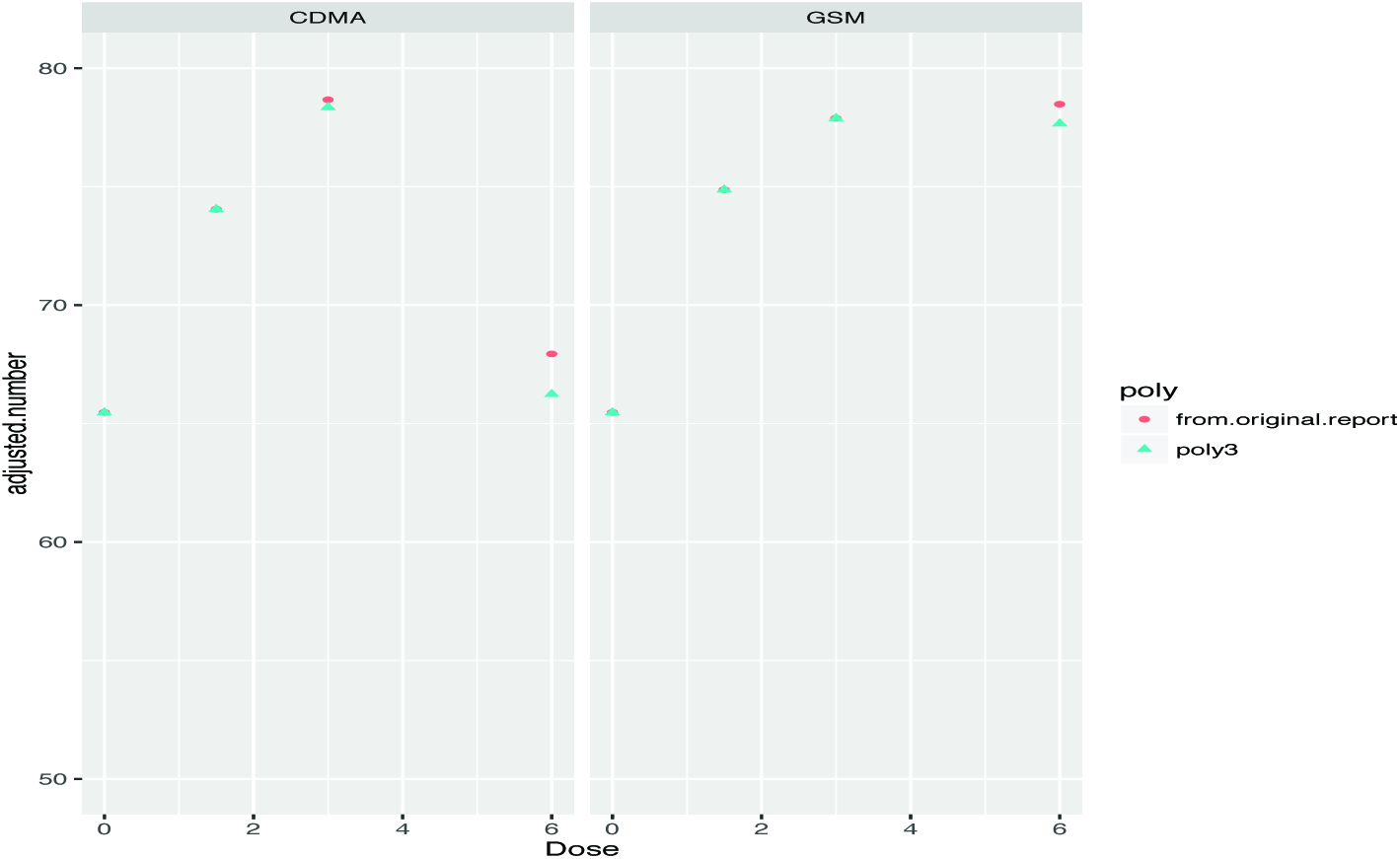
**Comparison of poly-3 adjusted rat numbers between those from the original report versus those calculated from the raw data**.

The poly-3 adjusted rat number from Table 3 of the original report is compared with the poly-3 adjusted rat number that I calculated from the raw data for heart schwannomas experiment.

**Figure S2.**
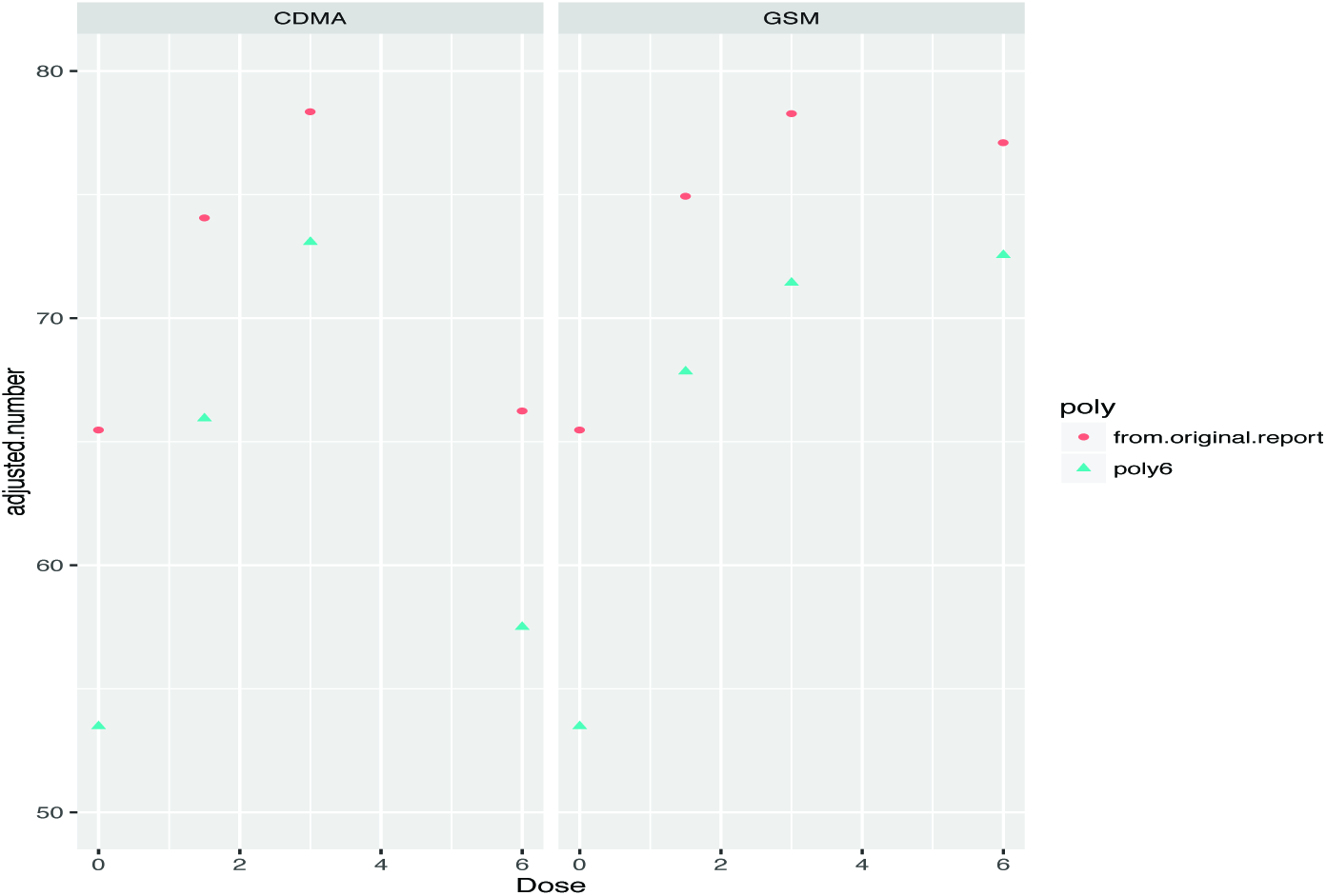
**Comparison of poly-6 adjusted rat numbers between those from the original report versus those calculated from the raw data**.

The poly-6 adjusted rat number from Table 1 of the original report is compared with the poly-6 adjusted rat number that I calculated from the raw data for brain gliomas experiment.

**Figure S3.**
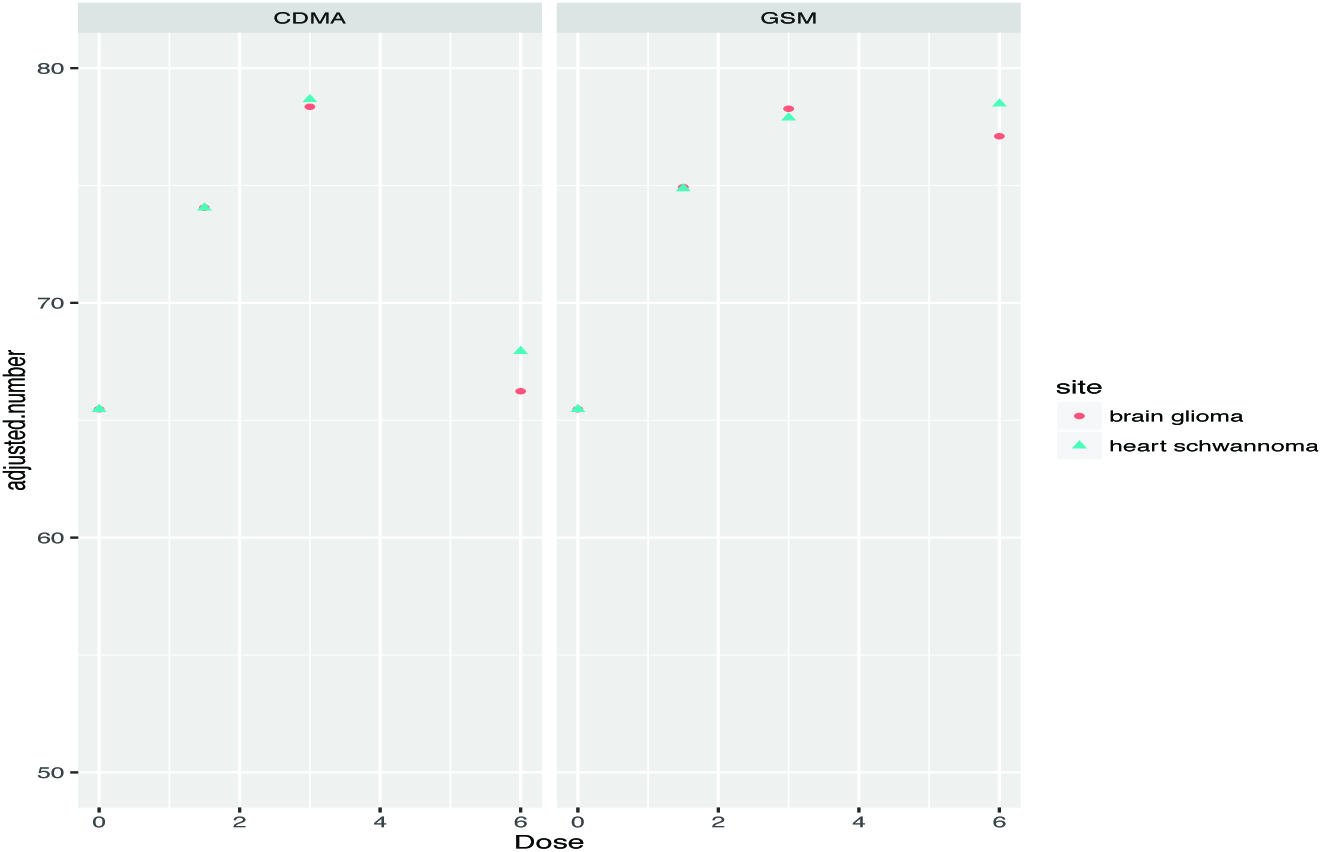
**Comparison of poly-6 adjusted rat numbers between those from the original report versus those calculated from the raw data**.

The adjusted rat numbers from Table 1 and Table 3 of the original report are compared with each other.

Reviewer: Aleksandra M. Michalowski, M.Sc., Ph.D., Laboratory of Cancer Biology and Genetics, NCI

**REVIEWER COMMENTS**

**Reviewer’s Name:**

Aleksandra M. Michalowski, Ph.D., M.Sc., National Cancer Institute/LCBG

**Report Title:**

Report of Partial Findings from the National Toxicology Program Carcinogenesis Studies of Cell Phone Radiofrequency Radiation (Whole Body Exposures); Draft 3-16-2016

**Charge:** To peer review the draft report and comment on whether the scientific evidence supports NTP’s conclusion(s) for the study findings.

1. Scientific criticisms:

a. *Please comment on whether the information presented in the draft report, including presentation of data in any tables, is clearly and objectively presented. Please suggest any improvements*. Overall, the information included in the report is presented in a comprehensive and accurate manner. Specifically, the experimental design and conditions are sufficiently documented and the choice of statistical approaches is explained; the results are well organized and necessary details are provided. Nevertheless, a few additions could be suggested:
  1. Appendix tables for all poly-k tests performed could be added. I believe this would enhance the presentation of the adjusted rates and the strength of the statistical evidence. As a possible example I prepared the below table using the R package *MCPAN* and its *poly3test(*) function.

  2. In the portion of the text describing poly-k test results, p-values are given for significant pairwise comparisons; I would also give the p-values estimated for the significant trends (maximum test).
  3. Information could be included regarding the software or programming environment used for the computations.
  4. In the portion of the text describing differences in survival at the end of the study between control and RFR-exposed animals (page 5§2) the compared characteristic is not named (median survival, TSAC?) and also no numerical values of the estimates or the range of differences are given. I would add numbers in the text or an Appendix table showing the group survival estimates described in this paragraph.

| poly-3 | Heart Schwannoma Malignant, Male |  |  |  | Heart Schwannoma Malignant, Female |  |  |  |
| --- | --- | --- | --- | --- | --- | --- | --- | --- |
| CDMA exposure | 0 | 1.5 | 3 | 6 | 0 | 1.5 | 3 | 6 |
| X | 0 | 2 | 3 | 6 | 0 | 2 | 0 | 2 |
| N | 90 | 90 | 90 | 90 | 90 | 90 | 90 | 90 |
| adjusted n | 63.8 | 72.4 | 77.1 | 66.6 | 67.9 | 71.8 | 70.3 | 78.0 |
| Dunnett contrast | — | 1.5 - 0 | 3 - 0 | 6 - 0 | — | 1.5 - 0 | 3 - 0 | 6 - 0 |
| Estimate | 0 | 0.03 | 0.04 | 0.09 | 0 | 0.03 | 0 | 0.03 |
| Statistic | — | 1.24 | 1.58 | 2.45 | — | 1.26 | 0 | 1.24 |
| p-value | — | 0.2704 | 0.1542 | **0.0209** | — | 0.2466 | 0.7992 | 0.2562 |
| Williams contrast | — | (6,3,1.5) - 0 | (6,3) - 0 | 6 - 0 | — | (6,3,1.5) - 0 | (6,3) - 0 | 6 - 0 |
| Estimate | 0 | 0.05 | 0.06 | 0.09 | 0 | 0.02 | 0.01 | 0.03 |
| Statistic | — | 2.78 | 2.75 | 2.45 | — | 1.27 | 0.88 | 1.24 |
| p-value | — | **0.0056** | **0.0060** | **0.0138** | — | 0.1661 | 0.2871 | 0.1744 |

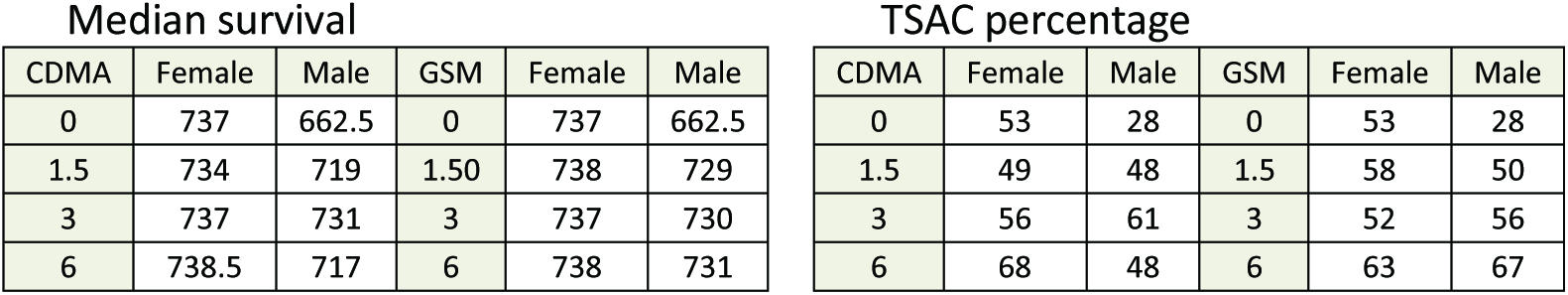
b. *Please comment on whether NTP’s scientific interpretations of the data are objective and reasonable. Please explain why or why not.* Appropriate statistical design and methods were applied in accord with the FDA/NTP guidelines for conducting long-term rodent carcinogenicity studies and analyses. The results and limiting issues were objectively discussed. The critical issue of shorter survival in the male control group was addressed with regard to the percentage of animals surviving to terminal sacrifice in historical control data (avg. 47%, range 24% to 72%) and the possible impact of the observed age of tumor occurrence on the statistical inference. I believe detailed information about animal selection and randomization procedures should be given so that the potential for allocation bias could be judged. As shown in the figure below, the lower survival rate to terminal sacrifice (28%) in the male control is accompanied by the higher rate of moribund sacrifice (49%); in the male group exposed to CDMA with 6 W/kg, a higher rate of natural death was observed (46%). It has been reported that insufficient randomization can lead to differences in survival rates. As an example, in a carcinogenicity study on aspartame it was suggested that lack of randomization to different rooms may have possibly been the cause of low survival rates (27%) in the control female group due to a high background infection rate (EFSA, 2006; Magnuson, B., Williams, G.M., 2008).

| Median survival |  |  |  |  |  | TSAC percentage |  |  |  |  |  |
| --- | --- | --- | --- | --- | --- | --- | --- | --- | --- | --- | --- |
| CDMA | Female | Male | GSM | Female | Male | CDMA | Female | Male | GSM | Female | Male |
| 0 | 737 | 662.5 | 0 | 737 | 662.5 | 0 | 53 | 28 | 0 | 53 | 28 |
| 1.5 | 734 | 719 | 1.50 | 738 | 729 | 1.5 | 49 | 48 | 1.5 | 58 | 50 |
| 3 | 737 | 731 | 3 | 737 | 730 | 3 | 56 | 61 | 3 | 52 | 56 |
| 6 | 738.5 | 717 | 6 | 738 | 731 | 6 | 68 | 48 | 6 | 63 | 67 |

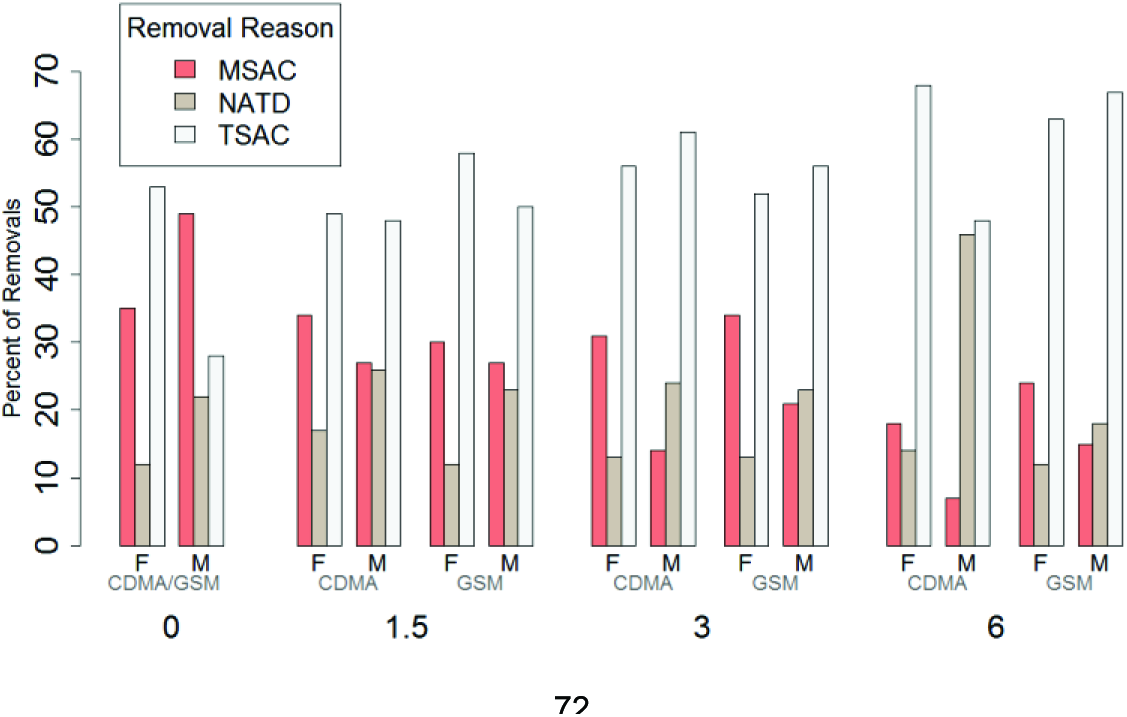
2. Please identify any information that should be added or deleted: A statement of the required statistical significance level should be added. FDA guidance suggests the use of significance levels of 0.025 and 0.005 for tests for positive trends in incidence rates of rare tumors and common tumors, respectively; for testing pairwise differences in tumor incidence the use of significance levels of 0.05 and 0.01 is recommended for rare and common tumors, respectively. If power calculations to determine the required sample size were performed, the results should also be included.
3. The scientific evidence supports NTP’s conclusion(s) for the study findings: *The NTP’s overall draft conclusion was as follows: “Under the conditions of these studies, the observed hyperplastic lesions and neoplasms outlined in this partial report are considered likely the result of exposures to test article A and test article B. The findings in the heart were statistically stronger than the findings in the brain.”* In my view, the results support the conclusion of likely carcinogenic effect of the RFR-exposure on Schwannoma heart lesions in male Harlan Sprague Dawley rats. Possible carcinogenic effects in the brain are marginal and are not sufficiently supported by statistical evidence in the male Harlan Sprague Dawley rats. In the female Harlan Sprague Dawley rats very few lesions were observed in either site and statistical significance was not reached at all.

Reviewer: R. Mark Simpson, D.V.M., Ph.D., Laboratory of Cancer Biology and Genetics, NCI

Analysis of National Toxicology Program (NTP) study evaluating risk in rat lifetime exposure to GSM or CDMA RFR.

Notes:

The NTP study document acknowledges several study limitations [page 10, discussion section]. Potential limitations should prominently factor into considerations regarding the context of the findings, as well as their interpretation and application.

<u>Working list of limitations potentially impacting NTP study interpretations</u>

- Difficulty in achieving diagnostic consensus in lesions classifications of rare, unusual, and incompletely understood lesion association
- Document appears to indicate that the second Pathology Working Group (PWG) empaneled to review and obtain lesion classification consensus, following the inability of the initial PWG to do so, may have reviewed different lesions sets
- No record of clinical disease manifestations due to lesions involving heart and brain [note lesions in heart and brain are mutually exclusive; affected rats have either one or the other and do not appear to have the involvement of both organs together (appendix E)]
- Lesions, including malignancies, do not appear to materially shorten lifespan, except for a subgroup of rats (less than 1/3 of affected rats) with malignant Schwannomas in heart
- Lack of shortened lifespan as a consequence of malignancy for the majority of affected rats contrasts with shortened lifespan of male control rats for which there is absence of attributable cause of death. The survival of the control group of male rats in the current study (28%) was relatively low compared to other recent NTP studies (avg 47%, range 24 to 72%). Creates greater reliance on statistical controlling for survival disparities and reliance on historical controls
- Reliance on historical controls made up of rats of different genetic strain background, held under different environmental conditions
- Absence of data on incidence of more frequently expected tumor occurrences in rats (background lesions)

Documenting the nature of the brain and cardiac lesions observed in RFR exposed rats and placing them into test article exposure-related context, in contrast to potential for their occurring spontaneously, are important and challenging goals. The NTP study limitations make the interpretation of reasonable risk more complicated. NTP acknowledgements of study limitations appear factored into one of NTP’s reviewer’s study conclusion, i.e., findings represent “some evidence” for a test article effect in statistically significant trend for Schwannomas; an opinion which is coupled with a conclusion for “equivocal evidence” of an effect in relation to malignant gliomas of the brain [NTP Appendix F, Reviewer Comments].

The summation from Appendix F reviewers regarding existence of test article effect is less than conclusive. The NTP study documents a series of cytoproliferative changes in heart and brain. The nature of some of the changes is challenging diagnostically and appears to be incompletely understood. These findings are presented in the absence of complete analysis of the entire consequences of the study effects. For example, no potential significance for test article effect context is given to any of granular cell proliferative lesions of the brain, a finding mentioned only as a contrast to what was less well understood pathologically (NTP Appendix C, Pathology). It is noteworthy that the lesion types analyzed in the NTP RFR study under review are uncommon historically in rats, in the organs discussed. Furthermore, the malignancies of neuroglia appear to be paired with the occurrence of poorly understood changes involving neuroglial cell hyperplasias in the central and peripheral nervous systems. Little information can be gleaned from the literature about the nature and significance of these latter proliferative changes, interpreted by NTP as nonneoplastic and non-inflammation-reactive neuroglial cell in nature. Although unclear in the NTP study document, it is plausible that the particular lesion constellation, along with the relative novelty of some lesions, contributed to the lack of consensus regarding the nature of the lesions on the part of the initial PWG study pathologists. Concern raised by one of the reviewers (Appendix F, Reviewer Comments) regarding how this difficulty in ability to classify lesions might impact comparisons to historical control lesion incidence data (NTP Table D) is certainly principled.

The extraordinary PWG process, presumably posed by the difficult diagnostic interpretations, has the potential to influence the reliance on historical controls. In this regard, study limitations concerning determination of whether or not there is a test article effect include the substantially poor survival of male rats in the control group. The survival of the control group of male rats in the study under review (28%) was relatively low compared to other recent NTP studies (avg 47%, range 24 to 72%). This apparently led to greater statistical construction to account for the impact of study matched controls, and created increased reliance upon historical data of rare tumor incidences in control animals taken from other chronic carcinogenicity studies. NTP acknowledges a limitation in using the historical incident data and a small study match control group due to poor survivability. There are potential sources of variability when using historical controls of different rat strains and fluctuating study conditions (environment, vehicle, route of exposure, etc.), as is the case here. It seems less than clear what appropriate background lesion incidence is, as NTP indicates some data involve other strains of rats. The range of lesion incidence in historical controls could mean that the true incidence of some lesions varies considerably and might be considered rare or more common depending upon the incidence rate.

The guidance manual on Statistical Aspects of the Design, Analysis and Interpretation of Chronic Rodent Carcinogenicity Studies of Pharmaceuticals by the FDA provided for this review discusses applying comparisons using historical control lesion incidences at some length [beginning page 27, line 996]. Considering lesions as being rare or more common appears to influence selection of the level of statistical significance for comparisons. It appears that analysis for significant differences in tumor incidence between the control and the dose groups for these NTP studies has been established at the 0.05 level (NTP Tables 1,3,5). Interpretations of trend tests may be influenced by the choice of decision rule applied. Such choices can result in about twice as large overall false positive error as that associated with control-high pairwise comparison tests [page 28, line 1012-1026]. The FDA guidance manual [page 31, line 1136] highlights concern regarding reliance upon historical control incidence data, stating that using historical control data in the interpretation of statistical test results is not very satisfactory because the range of historical control rates is usually too wide. This is especially true in situations in which the historical tumor rates of most studies used are clustered together, but a few other studies give rates far away from the cluster. When the range of historical control data is simply calculated as the difference between the maximum and the minimum of the historical control rates, the range does not consider the shape of the distribution of the rates. These circumstances may impose some limitations on optimal risk assessment designs.

Somewhat paradoxically then, NTP study limitations including that imposed due to reliance upon less than optimal historical control lesion incidence data for much of the comparisons between treated and untreated rats, is confronted by existence of a difficult to classify and incompletely understood lesion constellation interpreted to include neuroglial cell hyperplasia. Notwithstanding, this confounding proliferative lesion occurring in the context along with malignancies of apparently similar histogeneses, sustains a level of concern for a rare injury mechanism related to test article effect. Additional information about the study together with an assessment of the statistical analyses may enhance the value of this analysis.

R. Mark Simpson, D.V.M., Ph.D.

## Appendix G2: NTP’s Responses to NIH Reviewers’ Comments

Reviewers: R. Mark Simpson, D.V.M., Ph.D. and Diana Copeland Haines, D.V.M.

<u>*Responses Relating to the Pathology Review Process*</u>

> Drafts of the PWG reports are provided. As described in the PWG report, the specific task of the first PWG (January 29^th^, 2016) was to: 1) confirm the presence of glial cell hyperplasia and malignant gliomas in the brain and Schwann cell hyperplasia and schwannomas in the heart; 2) develop specific diagnostic criteria in the brain for distinguishing glial cell hyperplasia from malignant glioma and gliosis, and in the heart for distinguishing between Schwann cell hyperplasia and schwannoma. The PWG participants confirmed the malignant gliomas and schwannomas, but the criteria for distinguishing between hyperplasia and neoplasia differed between the participants.
>
> In order to clearly establish specific diagnostic criteria for the differentiation between hyperplastic and neoplastic lesions in the brain and heart, two additional PWGs were convened. The participants for the second (February 25, 2016) and third (March 3, 2016) PWGs were selected based on their distinguished expertise in the fields of neuropathology and cardiovascular pathology, respectively. Some of the participants were leaders in the International Harmonization of Nomenclature and Diagnostic Criteria initiative. The neuropathology experts of the second PWG confirmed the malignant gliomas in the brain, established diagnostic criteria for glial cell hyperplasia, and agreed that the hyperplastic lesions are within a continuum leading to malignant glioma. The cardiovascular pathology experts of the third PWG established specific diagnostic criteria for Schwann cell hyperplasia and schwannoma in the endocardium and myocardium, and reviewed and confirmed all cases of Schwann cell hyperplasia and schwannoma observed in these studies. The outcome of the PWG provided a very high degree of confidence in the diagnoses.
>
> The participants of the first PWG (January 29^th^, 2016) only reviewed a subset of the glial lesions that were observed in the studies. The review for the second PWG (February 25, 2016) included all glial lesions in the studies including the subset that was reviewed in the first PWG.

<u>*Responses Relating to Considerations of Historical Control Data*</u>

> For NTP toxicology and carcinogenicity studies, the concurrent controls are always the primary comparison group. However, historical control information is useful particularly in instances when there is differential survival between controls and exposed groups, as was observed in the RFR studies. Rates for glial cell neoplasms and heart schwannomas from control groups of male Harlan Sprague Dawley rats from other recently completed NTP studies are presented in Appendix D of the 3-16-2016 draft report. While Harlan Sprague Dawley rats are an outbred strain, they are considered a single genetic strain in the same sense as other outbred strains, such as the Long-Evans or Wistar rat. Therefore, these historical control tumor rates are applicable to this study. However, it’s important to note that the studies listed in Appendix D were carried out at laboratories other than the RFR studies, and under different housing and environmental conditions. At the time of the 3-16-2016 draft report, not all of these studies had undergone a complete pathology peer review. In the past several weeks NTP pathologists have reviewed brain and heart slides from these male rat control groups, and have confirmed, with few exceptions, the low rates of hyperplastic and neoplastic lesions reported in Appendix D, applying the diagnostic criteria established during the PWGs outlined in Appendix C.

*Given the multiple comparisons inherent in this kind of work, there is a high risk of false positive discoveries (Michael S. Lauer).*

> Although the NTP conducts statistical tests on multiple cancer endpoints in any given study, numerous authors have shown that the study-wide false positive rate does not greatly exceed 0.05 (Fears et al., 1977; Haseman, 1983; Office of Science and Technology Policy, 1985; Haseman, 1990; Haseman and Elwell, 1996; Lin and Rahman, 1998; Rahman and Lin, 2008; Kissling et al., 2014). One reason for this is that NTP’s carcinogenicity decisions are not based solely on statistics and in many instances statistically significant findings are not concluded to be due to the test agent. Many factors go into this determination including whether there were pre-neoplastic lesions, whether there was a dose-response relationship, biological plausibility, background rates and variability of the tumor, etc. Additionally, with rare tumors especially, the actual false positive rate of each individual test is well below 0.05, due to the discrete nature of the data, so the cumulative false positive rate from many such tests is less than a person would expect by multiplying 0.05 by the number of tests conducted (Fears et al., 1977; Haseman, 1983; Kissling et al., 2015).

***I’m getting slightly different values for poly-k adjusted denominators (Michael S. Lauer*)**.

***I compared poly–-3 adjusted number from Table 3 in the original report versus the poly–-3 adjusted number that I calculated using the raw data from the excel files. Supplementary Figure S1 shows that these two sets of numbers agree with each other in general. This is in contrast to the comparison for poly–6 adjusted number from Table 1 in the original report versus the poly–-6 adjusted number that I calculated using the raw data from the excel files (Supplementary Figure S2). In fact, the adjusted rat numbers from Table 1 and Table 3 of the original report look quite similar (Supplementary Figure S3). This suggests that the poly–-3 adjusted number was used in the footnotes in both Table 1 and Table 3 in the original report. (Max Lee*)**

***I noted that in Table S2 the adjusted numbers in from.original.report and poly3 are identical at Dose 0 and 1.5 for both CDMA and GSM as well as at Dose 3 for GSM but differ slightly in the other treatment doses for heart schwannomas. One possible cause of the difference is that the version of the raw data in the excel files differs from that used to generate the original report. The second possibility is typo in the footnote in Table 3. I also generated Table S3 that has the poly–-6 adjusted numbers for brain gliomas. The two sets of the poly–-6 adjusted numbers are very different. (Max Lee*)**

***Information could be included regarding the software or programming environment used for the computations. (Aleksandra M. Michalowski*)**

> The adjusted denominators in Table 1 of the original report were labeled as poly-6 denominators, but were actually poly-3 denominators. This error was noted and brought to Dr Tabak’s attention by Dr. Bucher in a March 22 email.
>
> The p-values and adjusted denominators calculated by NTP are correct, except as noted for Table 1, and were calculated using validated poly-k software. This software is coded in Java and is embedded within NTP’s TDMSE (Toxicology Data Management System Enterprise) system. Poly-k calculations conducted by the reviewers in R may vary slightly from the NTP’s calculation due to selection of study length and the NTP’s use of the Bieler-Williams variance adjustment and a continuity correction. In his calculations, Dr. Lauer used 90 weeks as the study length, whereas the actual study length was 104 weeks. It is not apparent from the R documentation that the Bieler-Williams adjustment or the continuity correction is incorporated into the poly-3 calculations in R. In his calculations, Dr. Lee used two-sided p-values. In NTP statistical tests for carcinogenicity, the expectation is that if the test article is carcinogenic, tumor rates should increase with increasing exposure; thus, the NTP employs one-sided tests and p-values are one-sided. Using one-sided p-values in Dr. Lee’s Table 1, the GSM trend if there were 1 brain glioma in the control group remains nonsignificant, but the CDMA trend approaches 0.05 (p = 0.054) if there were 1 brain glioma in the control group. In Dr. Lee’s Table 2, the one-sided p-value for the GSM trend if there were 1 heart schwannoma in the control group approaches 0.05 (p = 0.054) and the one-sided p-value for the CDMA trend in heart schwannomas remains significant at p = 0.018 if there were 1 heart schwannoma in the control group. In Dr. Lee’s Table 3, the one-sided p-value for the CDMA pairwise comparison is significant at p = 0.049 if there were 1 heart schwannoma in the control group.

***A statement of the required statistical significance level should be added. FDA guidance suggests the use of significance levels of 0.025 and 0.005 for tests for positive trends in incidence rates of rare tumors and common tumors, respectively; for testing pairwise differences in tumor incidence the use of significance levels of 0.05 and 0.01 is recommended for rare and common tumors, respectively. (Aleksandra M. Michalowski*)**

> Although the FDA guidance suggests lowering the significance level for most tests of trend and pairwise differences, this guidance is based on a misunderstanding of findings reported by Haseman (1983). In this paper, Haseman discusses several rules proposed by others for setting the significance level lower than 0.05. <u>*Iƒ*</u> these rules are rigidly followed, Haseman showed that study conclusions will be consistent with the NTP’s more complex decision-making process, for which 0.05 is the nominal significance level and p-values are taken into consideration along with other factors (outlined above in response to comment 1) in determining whether the tumor increase is biologically significant. The NTP does not strictly adhere to a specific statistical significance level in determining whether a carcinogenic effect is present.

***Appendix tables for all poly-k tests performed could be added. (Aleksandra M. Michalowski*)**

> Dr. Michalowski proposed a sample table. The rows corresponding to X, N, adjusted n are already included in the tables or appear the footnotes in the tables. The rows corresponding to “Dunnett contrast” and “Williams contrast” are not appropriate for dichotomous tumor data. Both Dunnett’s test and Williams’ test assume that the data are continuous and normally distributed.

***In the portion of the text describing poly-k test results, p-values are given for significant pairwise comparisons; I would also give the p-values estimated for the significant trends. (Aleksandra M. Michalowski*)**

> Indicators of significant trends are given in the tables in the form of asterisks next to control group tumor counts.

***There are a couple of errors in the footnote of Table 3 in the original report. 2/74.05 (5%) should be 2/74.05 (2.7%). 3/78.67 (4%) should be 3/78.67 (3.8%). (Max Lee*)**

> Thank you for pointing this out. The percentages will be corrected in our final report.

***Were control rats selected in utero like the exposed rats were? Were pregnant dams assigned to different groups by formal randomization ? How were the 3 pups per litter chosen ? (Michael S. Lauer).***

***I believe detailed information about animal selection and randomization procedures should be given so that the potential for allocation bias could be judged. (Aleksandra M. Michalowski*)**

> Pregnant dams were assigned to groups, including the control group, using formal randomization that sought to also equalize mean body weights across groups. The three pups per sex per litter were selected using formal randomization, as well. Tumors in the heart and brain were not observed in littermates, indicating that there was no litter-based bias in the results.

***Were all analyses based on the intent-to-treat principle? Were there any crossovers? Were all rats accounted for by the end of the experiment and were all rats who started in the experiment included in the final analyses? (Michael S. Lauer*)**

> The intent-to-treat principle is not relevant to this animal experiment, in which all animals that were assigned to a treatment group received the full and equal treatment of that group. There were no crossovers. All animals that started the experiment were accounted for by the end of the experiment and included in the final analyses.

***The PWG review blinding was not complete. (Michael S. Lauer*)**

> PWG reviewers were blinded to the identity of the test article and the level of exposure but were not blinded to the fact that there were two different, yet related, test articles (modulations of cell phone RFR), to emphasize the fact that there was a common control group.

***Did the authors perform a prospective sample size calculation ? (Michael S. Lauer*)**

***If power calculations to determine the required sample size were performed, the results should also be included. (Aleksandra M. Michalowski*)**

> Sample size calculations were conducted for this study. However, for detecting carcinogenesis, sample size and power will depend on the baseline (control) tumor rate and the expected magnitude of the increase in tumors. For example, at 80% power, sample size requirements will be quite different for detecting a 2-fold increase in a rare tumor having a spontaneous occurrence of 0.5% compared to a 2-fold increase in a more common tumor having a spontaneous occurrence of 10%. Because many different tumor types having a wide range of spontaneous occurrence are involved in these studies, there is no “one-size-fits-all” sample size; rather, the sample size is a compromise among several factors, including obtaining reasonable power to detect moderate to large increases for most tumor types, while staying within budgets of time, space, and funding. A sample of 90 animals per sex per group was selected as providing as much statistical power as possible across the spectrum of tumors, under the constraints imposed by the exposure system.
>
> The NTP’s carcinogenicity studies are similar in structure to the OECD’s 451 Guideline for carcinogenicity studies and the FDA’s guidance for rodent carcinogenicity studies of pharmaceuticals. These guidelines recommend at least 50 animals of each sex per group, but also mention that an increase in group size provides relatively little increase in statistical power. In the NTP’s RFR studies, the group sizes were 90 animals of each sex per group, nearly twice as many as the minimum recommendation. Increasing the group sizes further provides diminishing returns, for which additional animals do not substantially increase power.

***The low power implies that there is a high risk of false positive findings (citing Ioannidis, 2005). … I suspect that this experiment is substantially underpowered and that the few positive results found reflect false positive findings (citing Ioannidis, 2005). (Michael S. Lauer*)**

> It is true that the power is low for detecting moderate increases above a low background tumor rate of approximately 1 − 2 %, as was seen in the brain and heart tumors. However, this low power <u>does not</u> correspond to a high risk of false positive findings. The paper by Ioannidis that was cited correctly states that when studies are small or effect sizes are small (i.e., statistical power is low), “the less likely the research findings are to be true.” Research findings can be “not true” if the result is a false positive or a false negative. With low statistical power, false negatives are much more likely than false positives. Therefore, the vast majority of false research findings in a low power situation will result from the failure to detect an effect when it exists. The false positive rate on any properly constructed statistical test will not exceed its significance level, alpha. By definition, the significance level of a statistical test <u>is</u> its false positive rate, and it is typically selected by the researcher, often at a low fixed value such as 0.05 or 5%.

***If we were repeating the experiment, we may see some control studies have 1 or more tumors. (Max Lee) (Dr. Lee also presented analyses of the male rat data, inserting hypothetical data on one tumor-bearing animal in the control group.*)**

> In light of the historical control data, Dr. Lee demonstrated that several associations became less or not significant with the insertion of a tumor data point in the control group. While we appreciate that some other studies had one or more tumors, the NTP considers the concurrent control group as the most important comparator to the treated groups. We took the historical control tumor rates into account in a more subjective manner in our interpretation of the findings. In 2010, we asked to adopt a more formal method of incorporating historical control data in our statistical testing, but our Board of Scientific Counselors voted against adopting the method.

***It is puzzling why the control had short survival rate. Given that most of the gliomas and heart schwannomas are late-developing tumors, it is possible that if the controls were living longer some tumors might develop. Although the use of poly-3 (or poly-6) test intended to adjust the number of rats used in the study, it is still important to re-evaluate the analysis by considering the incidence rate in controls not being 0. (Max Lee*)**

> We do not know why the male rat control group had a low survival rate. We generally do observe lower survival rates in studies such as the RFR studies in which animals are singly-rather than group housed. While some tumors might possibly have arisen in controls if they lived longer, it was notable that no glial cell or Schwann cell hyperplasias were found in these animals as well.
>
> The poly-k (e.g., poly-3 or poly-6) test was developed to adjust for the fact that not all animals survive to the end of a two-year study, and survival rates may differ among groups. The test is essentially a Cochran-Armitage trend test in which the denominator of the tumor rate in each group is adjusted downward to better reflect the number of animal-years at risk during the study. Each animal that develops the tumor or survives to the end of the study is counted as one animal. Each animal that does not develop the tumor and dies (or is moribund sacrificed) before the end of the study is counted as a fractional animal. The fraction is calculated as the proportion of the study that it survived, raised to the k-th power; k = 3 or k = 6 in this study. The survival-adjusted tumor rate in each group is then the number of animals having the tumor of interest divided by the total count of animals at risk of developing the tumor in the group. These survival-adjusted rates are used in the Cochran-Armitage formula to provide the poly-k test for dose-related trends and pairwise comparisons with the control group.
>
> The poly-k test has been shown to yield valid inferences about tumor rates in NTP two-year rat and mouse carcinogenicity studies (Bailer and Portier, 1988; Portier and Bailer, 1989; Portier et al., 1986). Its theoretical basis is that tumor incidence, while not directly observed unless the tumor is immediately lethal, follows a Weibull distribution with a shape parameter, k. Verification using NTP studies has shown that if k is between 1 and 5, setting k = 3 yields a valid statistical test (Portier and Bailer, 1989; Portier et al, 1986). Thus, most of the time, the NTP uses the poly-3 test. If a tumor type is late-occurring, as we observed with the brain gliomas, k = 6 is a better fit to the data and the poly-6 test has more validity.

***In the portion of the text describing differences in survival at the end of the study between control and RFR-exposed animals the compared characteristic is not named and also no numerical values of the estimates or the range of differences are given. I would add numbers in the text of an Appendix table showing the group survival estimates described in this paragraph. (Aleksandra M. Michalowski*)**

> The Statistical Methods section describes the method for comparing survival distributions between the control and RFR-exposed groups, namely, Tarone’s (1975) life table test to identify exposure-related trends in survival and Cox’s (1972) method for testing two groups for equality of survival distributions.

## ADDITIONAL RESPONSE

Dear All,

Thanks again for all your helpful comments on the NTP RFR studies. I did want to follow up on one remaining point of disagreement that Mike Lauer alluded to in his comments about low powered studies. Although we agree that our study design had low power to detect statistically significant neoplastic effects in the brain and heart, which occurred with both RFR modulations in male rats, we disagree over the assertion that low power in and of itself, creates false positive results. We cited a handful of publications outlining the statistical arguments against this with specific respect to the NTP rodent cancer study design in our response to comments document sent earlier. Although Mike referred to the example of positive findings in underpowered epidemiology studies that could not be replicated in larger follow up studies, there is a growing literature alluding to this problem with respect to experimental animal studies as well. An example is a relatively recent article by one of our collaborators in CAMARADES, Malcolm MacLeod.

http://www.nature.com/news/2011/110928/full/477511a.html

It’s important to distinguish between low power to detect effects, and the constellation of other factors that often accompany low powered experimental animal studies in contributing to this problem. We’ve addressed this issue in a recent editorial, and these factors are captured in our published systematic review process for evaluating study quality in environmental health sciences (Rooney et al., 2014).

http://ehp.niehs.nih.gov/wp-content/uploads/122/7/ehp.1408671.pdf

http://ehp.niehs.nih.gov/wp-content/uploads/122/7/ehp.1307972.pdf

Table 1 in the Rooney et al. report outlines risk of bias considerations that commonly plague studies carried out by academic researchers that are accounted for in NTP studies.

I provide these examples to assure you that we are completely cognizant of these issues and take them very seriously. Again, we appreciate the help you’ve provided in assuring that we appropriately interpret and communicate our findings.

Best

John Bucher

## Footnotes

1 NTP is a federal, interagency program, headquartered at the National Institute of Environmental Health Sciences, part of the National Institutes of Health, whose goal is to safeguard the public by identifying substances in the environment that may affect human health. For more information about NTP and its programs, visit http://ntp.niehs.nih.gov

2 IARC (International Agency for Research on Cancer). 2013. Non-Ionizing Radiation, Part 2: Radiofrequency Electromagnetic Fields. IARC Monogr Eval Carcinog Risk Hum 102. Available: http://monographs.iarc.fr/ENG/Monographs/vol102/mono102.pdf [accessed 26 May 2016].

3 Specifications for the Conduct of NTP Studies, http://ntp.niehs.nih.gov/ntp/test_info/finalntp_toxcarspecsjan2011.pdf

* The term “carcinogenic” in this case refers to NTP studies in which any group of male or female rats or mice was judged to show “clear” or “some” evidence of carcinogenic activity. Keep in mind that many of these agents were selected for cancer studies based on a suspicion that they would cause cancer. Other agents, such as cell phone RFR, were chosen based more on the sheer numbers of people exposed. The “level of evidence” definitions are indicated below. As one can see, statistical significance is only one of many considerations that go into the study interpretation.

1 Pathology peer review of remaining lesions from the cell phone RFR studies continues and is not addressed in thisreport.

2 Locking data refers to restricting access to the computer database so the data for a particular study cannot be changed.

